# Single Cell Analysis Reveals Multi-faceted miR-375 Regulation of the Intestinal Crypt

**DOI:** 10.1101/2020.10.01.321612

**Authors:** Michael T. Shanahan, Matt Kanke, Ajeet P. Singh, Jonathan W. Villanueva, Adrian J. McNairn, Oyebola O. Oyesola, Alessandro Bonfini, Yu-Han Hung, Breanna Sheahan, Jordana C. Bloom, Rebecca L. Cubitt, Ennessa G. Curry, Wendy A. Pitman, Vera D. Rinaldi, Christopher M. Dekaney, Shengli Ding, Bailey C.E. Peck, John C. Schimenti, Lukas E. Dow, Nicolas Buchon, Elia D. Tait-Wojno, Praveen Sethupathy

## Abstract

The role of individual miRNAs in small intestinal (SI) epithelial homeostasis is under-explored. In this study, we discovered that miR-375 is among the most enriched miRNAs in intestinal crypts and stem cells (ISCs), especially facultative ISCs. We then showed by multiple manipulations, including CRISPR/Cas9 editing, that miR-375 is strongly suppressed by Wnt-signaling. Single-cell RNA-seq analysis of SI crypt-enriched cells from miR-375 knockout (375-KO) mice revealed elevated numbers of tuft cells and increased expression of pro-proliferative genes in ISCs. Accordingly, the genetic loss of miR-375 promoted resistance to helminth infection and enhanced the regenerative response to irradiation. The conserved effects of miR-375 were confirmed by gain-of-function studies in Drosophila midgut stem cells *in vivo.* Moreover, functional experiments in enteroids uncovered a regulatory relationship between miR-375 and Yap1 that controls cell survival. Finally, analysis of mouse model and clinical data revealed an inverse association between miR-375 levels and intestinal tumor development.

**Highlights:**

- miR-375 is one of the most enriched miRNAs in ISCs, especially facultative ISCs.
- miR-375 modifies tuft cell abundance and pro-proliferative gene expression in ISCs.
- Loss of miR-375 in mice enhances the host response to helminth infection and crypt regeneration.
- Mouse and human intestinal cancer are associated with reduced miR-375 expression.

**eTOC Blurb:** Sethupathy and colleagues show that miR-375 is a Wnt-responsive, ISC-enriched miRNA that serves as a break on intestinal crypt proliferation. They also show that miR-375 modulates tuft cell abundance and pro-proliferative gene expression in ISCs, that miR-375 loss enhances the host response to helminth infection as well as crypt regeneration post-irradiation, and its reduced expression is associated with intestinal cancer.

## Introduction

The small intestinal epithelium is a highly proliferative and regenerative tissue compartment of the intestinal mucosa. Actively-cycling intestinal stem cells (ISCs) located at the base of the epithelial crypts drive homeostatic renewal (3-5 days) of the epithelial lining (Barker, 2014), whereas facultative stem cells are induced upon injury to regenerate the epithelium (Ayyaz et al., 2019; Richmond et al., 2016). ISCs give rise to several different specialized cell types, which carry out diverse functions, including nutrient absorption and protection against pathogenic infection. To ensure the integrity of the epithelial lining and maintain proper cell lineage allocation, the proliferative capacity of actively-cycling ISCs, as well as injury-inducible facultative ISCs, must be tightly regulated by various cell-autonomous signaling pathways (Henning and von Furstenberg, 2016).

The Wnt signaling pathway is especially critical for determining ISC self-renewal and proliferative capacity (He et al., 2004). In the stem cells of other organ systems, regulatory RNAs (such as microRNAs) have been shown to play significant roles in modulating Wnt and other related signal transduction pathways (Peng et al., 2016; Zhang et al., 2019). However, the roles of most regulatory RNAs in the control of intestinal crypt behavior remain to be investigated.

MicroRNAs (miRNAs) are small, ~22nt regulatory RNAs that serve as fine-tuners of gene expression at the post-transcriptional level (Bartel, 2018; Gebert and MacRae, 2019). Their activity is critical in a wide array of biological processes, including growth and differentiation (Ivey and Srivastava, 2010). Over the past several years, it has been increasingly recognized that miRNAs impact the structure and function of the small intestine. McKenna and colleagues showed that gut-specific deletion of Dicer1, a critical processing enzyme of miRNAs, produces profound effects on intestinal architecture and allocation of mature cell types (McKenna et al., 2010). We previously reported that miRNAs in Sox9-Low jejunal epithelial cells, which are partially enriched for ISCs, are highly sensitive to microbes (Peck et al., 2017a). We also recently reported on a specific miRNA, miR-7, which is enriched along the enteroendocrine lineage trajectory and regulates intestinal epithelial proliferation (Singh et al., 2020). Others have shown that microRNAs such as miR-31, miR-34a, and miR-34b/c regulate mouse ISC proliferation (Bu et al., 2016; Jiang and Hermeking, 2017; Tian et al., 2017). Moreover, the evolutionary conserved role of microRNAs in regulating ISC activity is observed in the ability of let-7, miR-305, and miR-263a to modulate ISC division in *Drosophila melanogaster* (Chen et al., 2015; Foronda et al., 2014; Kim et al., 2017). Despite these advances, which miRNAs are enriched in mouse actively-cycling or facultative ISCs, and if/how they alter the cellular landscape and function of the intestinal crypt, remains unknown.

In this study, using several different reporter mice and fluorescence activated cell sorting (FACS) methods, we identify miR-375 as the most enriched miRNA in crypts (and especially facultative ISCs) relative to villus cells. Single cell RNA-seq reveals that genetic loss of miR-375 results in the elevation of Wnt signaling within ISCs and an increase in the number of tuft cells. Accordingly, we show that 375-KO mice exhibit significant reduction in worm burden after *Heligmosomoides polygyrus* infection and in the regenerative response to whole-body irradiation. The effects of miR-375 on intestinal epithelial survival and proliferation are further confirmed by functional experiments in *Drosophila* midgut stem cells *in vivo* and murine enteroids *ex vivo,* the latter of which also reveal a regulatory relationship between miR-375 and *Yap1.* Finally, analysis of data generated from mouse models and human samples reveals an association between miR-375 levels and intestinal tumor development.

## Results

### MiR-375 is the most enriched miRNA in intestinal stem cells

ISCs, transit-amplifying progenitors, and cells that contribute to the stem cell niche are enriched in intestinal crypts relative to villi (Barker, 2014). First, we profiled miRNAs, by small RNA sequencing (small RNA-seq), in mouse jejunal crypts and villi (fractions validated by RT-qPCR, **Fig. S1A,B**) separately and identified seven miRNAs that are significantly enriched in the crypts (fold change > 2.5, P-value < 0.05) (**Fig. 1A, Table S1**). To determine the extent to which ISCs contribute to the apparent crypt enrichment of these seven miRNAs, we performed small RNA-seq on three different sorted populations of crypt-resident cells that have previously been shown to exhibit stem cell features: the crypt-based columnar Lgr5-High cells (Barker et al., 2007) (validated by RT-qPCR, **Fig. S1C**), Sox9-Low cells (Formeister et al., 2009) (validated by RT-qPCR, **Fig. S1D**), and Cd24-Low cells (von Furstenberg et al., 2011). Bioinformatic analysis of the small RNA-seq data revealed that 14 miRNAs are shared among the top 20 most highly expressed miRNAs in each of the three cell populations (**Fig. 1B, Table S2**). Among these 14, only miR-375 is also among the seven crypt-enriched miRNAs (**Fig. 1A, Table S1**), which we further validated by quantitative real time qPCR (RT-qPCR) (**Fig. 1C**). Moreover, miR-375 is the most highly enriched miRNA in crypt-resident Lgr5-High ISCs compared to cells over-represented in villi (Sox9-Neg) (**Fig. 1D**). It is also the most highly enriched in crypt-resident Sox9-Low ISCs relative to Sox9-Neg (**Fig. 1E**). Other crypt-based cells include slowly-cycling Lgr5-negative facultative ISCs, (lower side population; LSP) (Dekaney et al., 2005) (validated by RT-qPCR, **Fig. S1E**). Although these cells are more slowly cycling than Lgr5+ ISCs at baseline, they have the capacity to revert to a more proliferative state in response to injury (Ayyaz et al., 2019; Richmond et al., 2016). We sorted and sequenced LSP cells and found that miR-375 is even more enriched in this population relative to Lgr5-High ISCs and upper side population (USP) actively-cycling stem cells (**Fig. 1F**), respectively. Taken together, these data indicate that miR-375 is over-represented in crypts, expressed in actively-cycling ISCs, and even more highly expressed in facultative ISCs.

**Figure 1.**
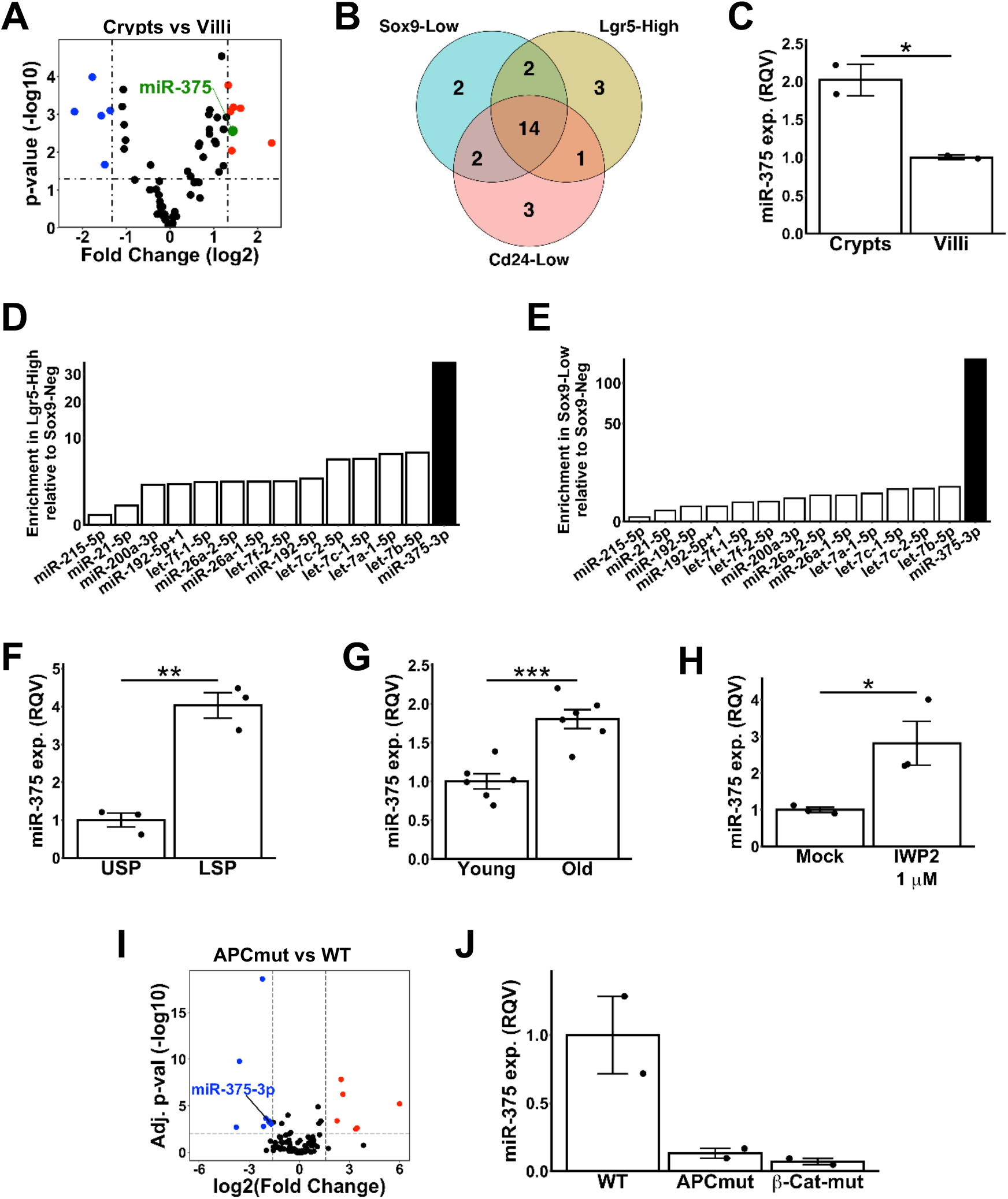
miR-375 is highly enriched in Lgr5-High ISCs and stem-like LSP cells. **A**, Volcano plot of differentially expressed microRNAs assessed by small RNA-seq analysis of isolated jejunal crypts relative to isolated jejunal villi from 3-5 months old C57BL/6 male mice (n=4). Only microRNAs with an RPMMM > 1000 in either crypt or villi were included in the analysis. Significant (P < 0.05; Student’s t-test) microRNAs with fold changes above 2.5 in intestinal crypts are colored red, while significant microRNAs with fold changes below −2.5 in intestinal crypts are colored blue. miR-375 is colored green. **B**, Venn diagram of the top 20 expressed microRNAs in Sox9-Low (blue), Lgr5-High (green), and Cd24-Low (red) FACS sorted cells. **C**, RT-qPCR data of miR-375 expression in jejunal crypts (n=2) of C57BL/6 mice relative to jejunal villi (n=2). **D**, Enrichment of the 14 microRNAs common to stem cell fractions in Lgr5-High cells versus Sox9-Neg cells. miR-375-3p is identified in black. **E**, Enrichment of the 14 microRNAs common to stem cell fractions in Sox9-Low cells versus Sox9-Neg cells. miR-375-3p is identified in black. **F**, RT-qPCR data of miR-375 expression of LSP sorted cells (n=3) relative to USP sorted cells (n=3). **G**, RT-qPCR data of miR-375 expression in jejunal crypts of 11-17 months old C57BL/6 male mice (Old, n=6) relative to 2-5 months old mice (Young, n=6). **H**, RT-qPCR data of miR-375 expression in jejunal enteroids established from tissue of WT B62J mice treated with 1 μM of Wnt antagonist IWP2 (n=3) relative to the mock condition (n=3). Representative of three independent experiments. **I**, Volcano plot of differentially expressed microRNAs assessed by small RNA-seq analysis of mouse jejunal enteroids with a Wnt activating APC mutation (APCmut) relative to non-mutated C57BL/6 wildtype (WT). Only microRNAs with a normalized count > 1000 in either WT or APCmut were included in the analysis. Significant (adjusted P < 0.01; DESeq2) microRNAs with fold change above 3 in APCmut enteroids are colored red, while significant microRNAs with fold change below −3 in APCmut enteroids are colored blue. miR-375 is identified in blue. **J**, RT-qPCR data of miR-375 expression in mouse jejunal enteroids with a Wnt activating APC mutation (APCmut, n=2) or a Wnt activating β-catenin mutation (β-Cat-mut, n=2) relative to non-mutated C57BL6/J (WT, n=2). * P < 0.05, ** P < 0.01, *** P < 0.001 by two-tailed Student’s t-test. RQV, relative quantitative value. RPMMM, reads per million miRNAs mapped.

### miR-375 expression is responsive to Wnt signaling in intestinal epithelial cells

Wnt signaling plays a prominent role in maintaining ISC survival and proliferative capacity (Flanagan et al., 2018). The high expression of miR-375 in Lgr5+ ISCs suggests that either miR-375 promotes or serves as a break on Wnt signaling pathways. Because miR-375 expression is even higher in facultative ISCs (**Fig. 1F**), which exhibit reduced Wnt signaling activity relative to Lgr5+ ISCs (Tao et al., 2015), we hypothesize that miR-375 and Wnt signaling are mutually suppressive.

To determine whether miR-375 is suppressed by Wnt signaling, we assessed miR-375 expression under various conditions in which Wnt signaling is perturbed. First, we considered the physiological condition of aging in which Wnt signaling in ISCs is reduced (Nalapareddy et al., 2017). Comparing expression in intestinal crypts of >1-year old mice relative to those derived from mice 5 months or younger, we observed a nearly 2-fold increase in mature miR-375 levels (**Fig. 1G**). Second, exposure of mouse jejunal enteroids to 1 μM IWP2, a porcupine inhibitor and Wnt antagonist (Mo et al., 2013), significantly increased expression of miR-375 relative to the mock condition (**Fig. 1H**). Finally, we used CRISPR/Cas9 to generate enteroids with a Wnt-activating mutation in APC (APCmut) (Dow et al., 2015) and performed small RNA sequencing analysis. We found that miR-375 is among the few miRNAs that are significantly suppressed in APCmut enteroids relative to unmutated control enteroid cells (**Fig. 1I, Table S3**). This finding was validated by RT-qPCR in APCmut enteroids as well as enteroids with a Wnt-activating β-catenin mutation (β-Cat-mut), also generated by CRISPR/Cas9 (**Fig. 1J**). Taken together, the results of these analyses indicate that miR-375 expression is controlled by the Wnt signaling pathway.

### miR-375 deficiency enhances small intestinal tuft cell abundance and enhances resistance to Heligmosomoides polygyrus helminth infection

The enrichment of miR-375 expression in intestinal stem cells and its responsiveness to Wnt signaling suggested that miR-375 may play a role in affecting the allocation and/or proliferation of various intestinal crypt cell types. To test this, we performed single cell RNA-seq (scRNA-seq) on jejunal crypt cells from 12-month-old WT mice and mice with a germ-line deletion for miR-375 (375-KO) (see Methods & Materials). Whole-body disruption of the miR-375 gene in 375-KO mice relative to WT was confirmed by tail genomic DNA genotyping (**Fig. S2A**) and by RT-qPCR data of miR-375 expression in jejunal enteroids derived from 375-KO mice relative to WT (**Fig. S2B**). In our analysis of the scRNA-seq data, we were able to identify several distinct clusters of cell populations (**Fig. 2A**) and unambiguously identify each of them as a well-established intestinal epithelial cell type based on the expression of distinguishing marker genes (**Fig. 2B**) as defined in a seminal scRNA-seq study of intestinal crypts by Haber and coworkers (Haber et al., 2017). In terms of the relative abundance of different cell types, the most striking result was a robust increase in tuft cells in 375-KO compared to WT (**Fig. 2C**). Gene expression analysis confirmed that several tuft cell marker genes including *Pou2f3*, *Gfi1b*, *Ascl2, Dclk1*, *Trpm5,* and *Rgs13* are elevated in 375-KO mice relative to WT (**Fig. 2D**). One of the best-characterized functions of intestinal tuft cells is to mediate the host anti-helminth response (Gerbe et al., 2016). Therefore, we hypothesized that the loss of miR-375 may reduce infectious burden. To test this hypothesis, both WT and 375-KO mice were infected with the helminth *Heligmosomoides polygyrus.* After 14 days of inoculation, although 375-KO mice did not display increased numbers of total immune cells (**Fig. S3A**) within the mesenteric lymph node, they did exhibit significantly increased numbers of eosinophils (**Fig. 2E**), which is a well-established marker of a robust host response to helminth infection. Consistent with this result, we found that 375-KO mice exhibited a significant decrease in intestinal worm burden (**Fig. 2F**), indicative of enhanced resistance to helminth infection.

**Figure 2.**
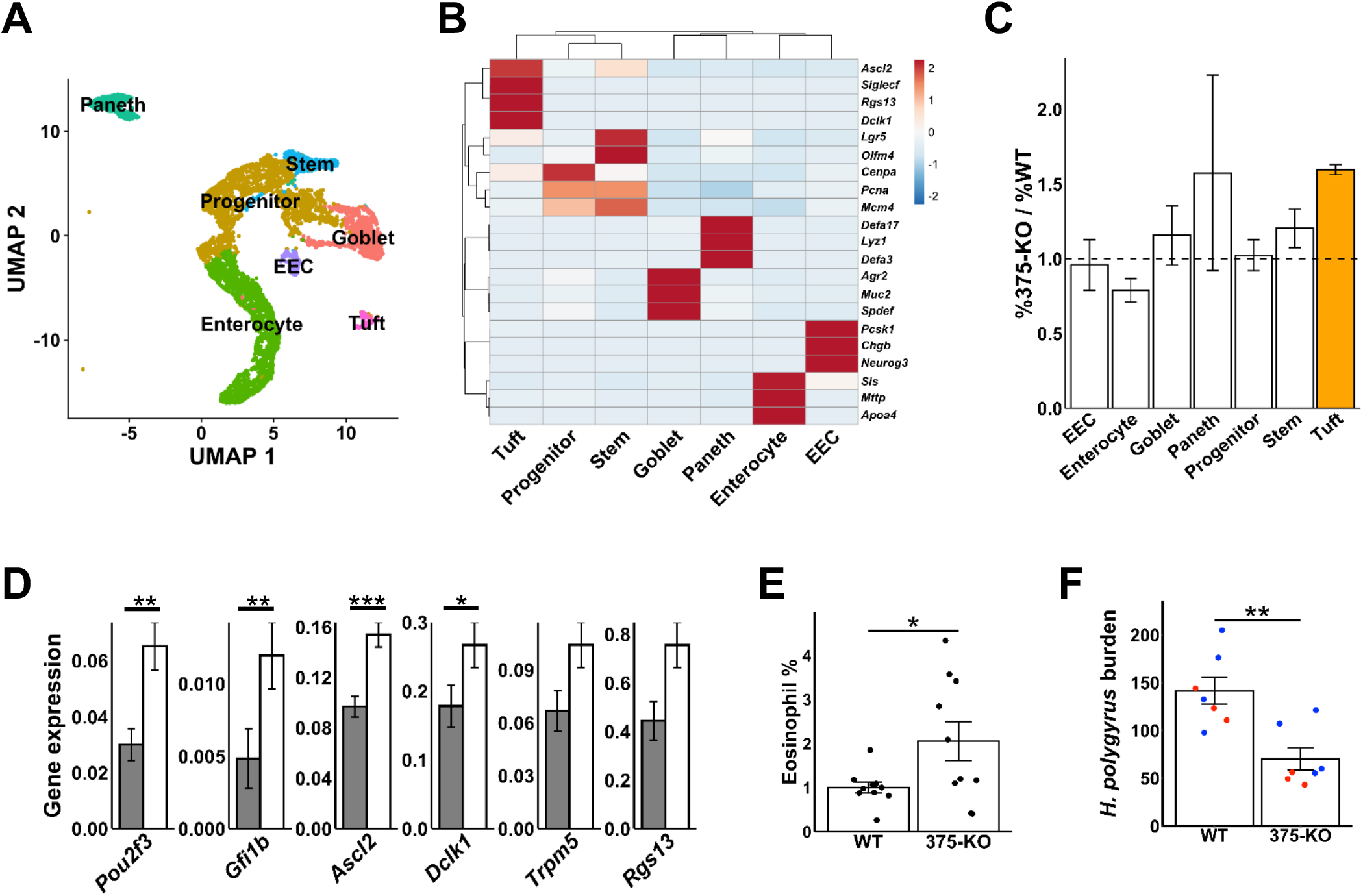
miR-375 deficiency increases small intestinal tuft cell numbers and enhances resistance to helminth *Heligmosomoides polygyrus* infection. **A**, UMAP scatterplot of 12 months old jejunal crypt cells from WT (n=2) and 375-KO (n=2) B62J mice analyzed by single cell RNA-seq. Cell clusters pertaining to the seven primary intestinal cell types are colored and labeled. **B**, Hierarchical clustering based on gene expression of classic markers of the seven small intestinal cell types. Red indicates enriched expression relative to all cells whereas as blue indicates de-enrichment of expression. **C**, Bar plot of the ratio of the percentage of 375-KO-derived cells within the various intestinal cell type clusters to the percentage of those derived from WT mice. **D**, Bar plot of transcript levels of tuft cell marker genes *(Pou2f3, Gfi1b, Ascl2, Dclk1, Trpm5,* and *Rgs13)* across all 375-KO jejunal crypt cells relative to WT. **E**, Bar plot of the percentage of eosinophils within isolated immune cell suspensions from FACS analyzed small intestinal tissue from WT (n-10) and 375-KO (n=10) mice after 14 days of infection with the helminth *Heligmosomoides polygyrus.* **F**, Bar plot of the number of *Heligmosomoides polygyrus* worms collected from the small intestinal lumen of WT (n=7) and 375-KO (n=7) mice after 14 days of infection (two representative experiments out of a total of three are shown; red – experiment #1, blue – experiment #2). * P < 0.05, ** P < 0.01, *** P < 0.001 by two-tailed Student’s t-test.

### miR-375 deficiency enhances Wnt signaling in intestinal stem cells and promotes the regenerative response to irradiation

Besides elevating tuft cell abundance, the scRNA-seq analysis of 375-KO and WT mice indicated that loss of miR-375 alters pathways in ISCs as well. Network analysis of the genes up-regulated in ISCs in 375-KO vs. WT showed enrichment of several transcription factors associated with the Wnt signaling pathway (**Fig. 3A**). Consistent with this result, we found that Wnt signaling target genes, including *Ccnd2, Cd44, Ascl2, Myc, Cldn2,* and *Trim65* are significantly up-regulated in ISCs from 375-KO mice compared to WT (**Fig. 3B,C**). These data suggested that miR-375 deficiency promotes Wnt signaling, which may elevate the proliferative capacity of the intestinal epithelium. The loss of miR-375 did not result in alteration of either mid-jejunal crypt depth (**Fig. S4A,B**) or the number of proliferating PH3+ mid-jejunal crypt cells (**Fig. S4C,D**) under unchallenged, baseline conditions. We hypothesized that the effect of miR-375 loss may be more pronounced in response to injury. To test this hypothesis, we evaluated the intestinal crypt response to whole-body irradiation in 375-KO mice compared to WT. Initial assessment of changes in mid-jejunal crypt depth of WT mice suggested that whole-body irradiation of 10 Gy results in peak crypt regeneration around 3 days post-irradiation (**Fig. S5A**). Therefore, we evaluated mid-jejunal crypt depth of WT and 375-KO mice during a period of increasing crypt regeneration at 1 day and 2.5 days post-irradiation (p.i.). Accelerated deepening of 375-KO mid-jejunal crypts in response to irradiation relative to WT indicated that reduced miR-375 expression potentiates intestinal regeneration induced by radiation injury (**Fig. 3D,E**). To determine whether other external challenges produce a similar phenotype, we subjected WT and 375-KO mice to a high fat diet for 16 weeks. There was no significant difference in weight gain between WT and 375-KO mice (**Fig. S4E**). Furthermore, both groups of mice manifested similar mid-jejunal crypt depths (**Fig. S4F,G**), indicating that the functional relevance of miR-375 to the regulation of crypt depth is specific to the regenerative response to irradiation and not generalizable to any stress condition.

**Figure 3.**
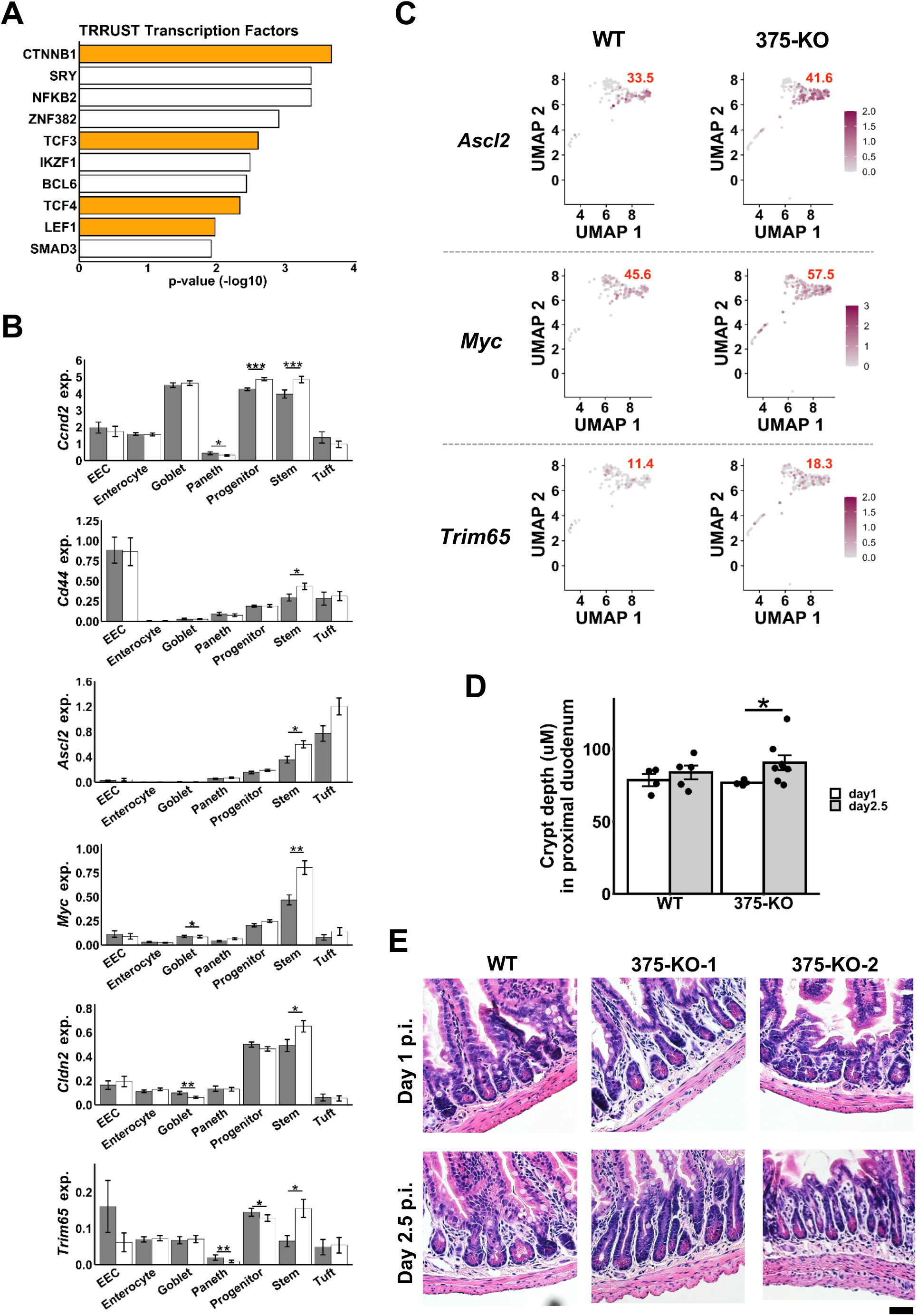
miR-375 deficiency augments the Wnt signaling pathway in ISCs and enhances the irradiation-induced regeneration response. **A**, Ranked bar plot of the −log10 P-values of the ten most enriched transcription factor regulators of the genes upregulated in the ISC cluster of 375-KO mice relative to WT mice. Established Wnt pathway transcription factors are denoted in orange. TRRUST is a manually-curated database of mouse and human transcriptional regulatory networks. **B**, Bar plots of the expression levels of Wnt signaling genes *Ccnd2, Cd44, Ascl2, Myc, Cldn2,* and *Trim65* within each of the cell clusters identified by scRNA-seq analysis in WT and 375-KO. **C**, UMAP scatter plots of a subset of genes shown in (B). Level of expression for the Wnt signaling genes *Ascl2*, *Myc*, and *Trim65* is shown in purple, while non-expressing cells are shown in grey. Only the ISC cluster is shown and the percentage of cells in the ISC cluster that are positive for the assayed gene is indicated in the top right corner in red. **D**, Bar plot of the mid-jejunal crypt depth for WT (n=4-5 mice) and 375-KO mice (n=4-7 mice) 1 day and 2.5 days after irradiation. **E**, Photomicrographs (x200) of H&E stained mid-jejunum in WT and 375-KO mice 1 day and 2.5 days post-irradiation (p.i.). Two different individual 375-KO mice are shown for 1 day and 2.5 days post-irradiation (p.i.). Yellow scalebar represents 50 μm. * P < 0.05, ** P < 0.01, ** P < 0.01 by one-tailed Student’s t-test.

### Loss of miR-375 leads to increased intestinal epithelial proliferation

We next sought to assess the function of miR-375 considering the intestinal epithelium alone. Specifically, we isolated jejunal crypts from 375-KO and WT mice and established enteroids *ex vivo.* The enteroids from 375-KO mice exhibited a significantly greater budding index relative to WT (**Fig. 4A,B**). 375-KO jejunal enteroids also displayed elevated survival compared with WT jejunal enteroids, an effect that was abrogated by inhibition of Wnt signaling (**Fig. 4C,D**).

**Figure 4.**
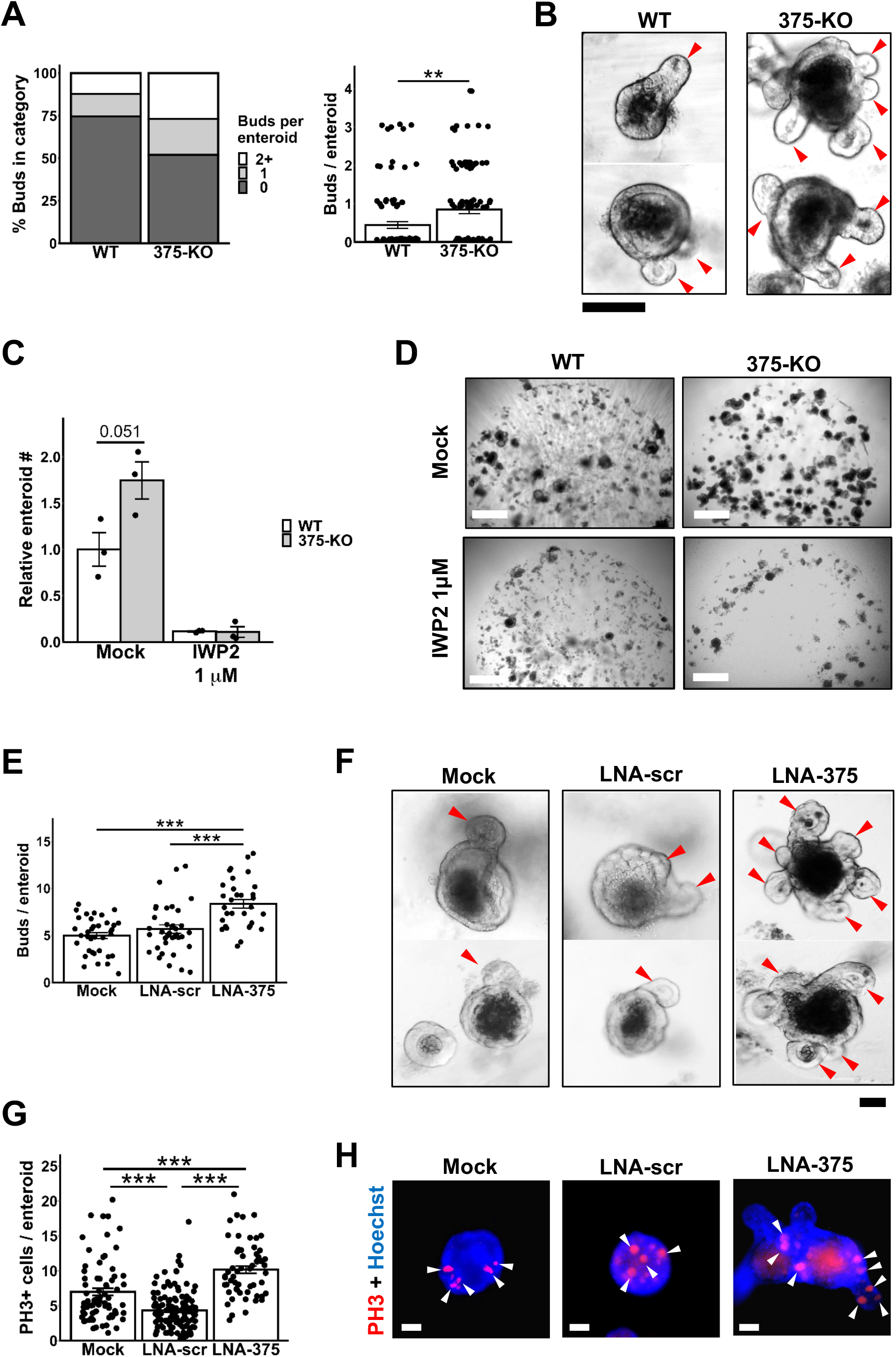
*In vivo* genetic loss and *ex vivo* knockdown of miR-375 promotes intestinal epithelial survival and proliferation. **A**, Plot of number of buds per enteroid of jejunal crypts derived from 375-KO relative to WT. **B**, Phase-contrast microphotographs (x400) of jejunal enteroids established from crypts of WT and 375-KO mice. Enteroid buds are indicated by red arrowheads. Yellow scalebar represents 50 μm. **C**, Plot of relative number of surviving jejunal enteroids from WT and 375-KO mice under conditions of mock treatment (n=3) or treatment with 1 μM of IWP2 (n=3). Values are expressed relative to mock treated WT mouse jejunal enteroids. Representative of three independent experiments (n=3-5 wells/group). **D**, Phase-contrast microphotographs (x50) of jejunal enteroids established from crypts of WT and 375-KO mice that were either mock treated or treated with 1 μM of IWP2. Yellow scalebar represents 500 μm. **E**, Plot of number of buds per enteroid of wildtype (WT) B62J mouse jejunal enteroids that were either under mock, locked nucleic acid (LNA)-scrambled (LNA-scr) or anti-miR-375 LNA (LNA-375) treated conditions (n=36 Mock, n=36 LNA-scr, n=31 LNA-375 enteroids examined). **F**, Phase-contrast microphotographs (x400) of WT mouse jejunal enteroids that were either mock, LNA-scr, or LNA-375 treated. Buds are indicated by red arrowheads. Yellow scalebar represents 50 μm. **G**, Plot of number of PH3+ cells per enteroid of mock, LNA-scr, and LNA-375 treated WT mouse jejunal enteroids (n=126 Mock, n=72 LNA-scr, n=58 LNA-375 enteroids examined). Yellow scalebar represents 50 μm. **H**, Fluorescent microphotographs (x400) of representative enteroids from WT mock, LNA-scr, and LNA-375 treated enteroid cultures. Hoechst-stained nuclei are shown in blue. PH3+ cells are shown in red and indicated by yellow arrowheads. Yellow scalebar represents 50 μm. **P< 0.01, *** P < 0.001 by two-tailed Student’s t-test.

We repeated the *ex vivo* study using enteroids from WT mice treated with either a locked nucleic acid (LNA) inhibitor of miR-375 (LNA-375) or LNA-scramble control (LNA-scr). Treatment with LNA-375 led to a ~1,000-fold loss in the levels of miR-375 (**Fig. S6A**). Though loss of miR-375 did not affect enteroid size (**Fig. S6B,C**), it did lead to significantly greater budding (**Fig. 4E,F**) and proliferation as measured by PH3+ cells/enteroid (**Fig. 4G,H**).

Given the extensive conservation of miR-375 across many species (**Fig. 5A**), we next evaluated the effect of miR-375 overexpression *in vivo* in the midgut epithelium of *Drosophila melanogaster,* a well-established model for gut stem cell response to stress and infection (Buchon et al., 2009). First, we performed overexpression of miR-375 using the lineage tracing system *esg^F/O^*. In this system, GFP and miR-375 are both expressed in esg+ ISCs and progenitor cells, as well as all direct progeny. We found that miR-375 reduces the generation of new GFP+ cells (**Fig. 5B**), suggesting that it suppresses ISC/progenitor cell activity. We next performed over-expression of miR-375 using *esg^TS^* (*esgGal4, Gal80^TS^>UAS-miR-375*), in which GFP and miR-375 are expressed only in esg+ ISCs and progenitor cells. We found that miR-375 did not alter the number of GFP+ ISCs and progenitor cells in basal homeostatic conditions (**Fig. 5C**). However, midguts under these basal conditions have a limited amount of proliferative activity; therefore, we repeated the experiment of miR-375 over-expression in ISCs and progenitor cells *(esgGal4, Gal80^TS^>UAS-miR-375*) under stress conditions of infection with *Erwinia carotovora carotovora* 15 (*Ecc*15), a fly pathogen that induces high levels of proliferation (Buchon et al., 2010). We found that miR-375 robustly decreases the number of GFP+ cells (**Fig. 5D**), suggestive of reduced cycling. Consistent with this finding, we also showed that over-expression of miR-375 in ISCs and progenitor cells led to a significant reduction in the total number of PH3+ proliferating cells (**Fig. 5E**). These results demonstrate that miR-375 is a strong cell-autonomous regulator of ISC proliferation and attest to the broad species conservation of the effect of miR-375 on intestinal epithelial proliferation.

**Figure 5.**
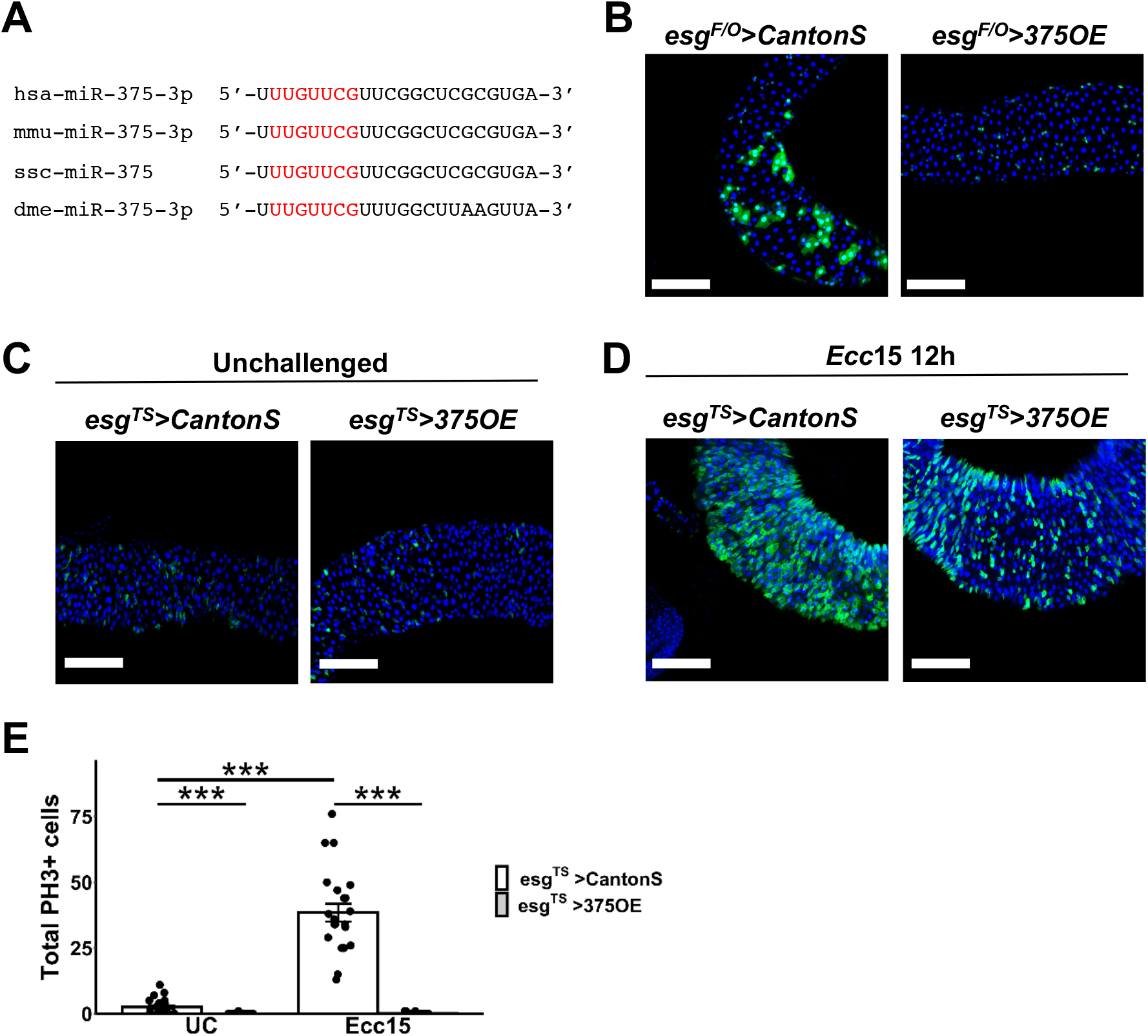
*In vivo* overexpression of miR-375 in Drosophila midgut ISCs reduces proliferation. **A**, Sequence alignment of mature miR-375 for *Homo sapiens* (hsa-miR-375-3p), *Mus musculus* (mmu-miR-375-3p), *Sus scrofa* (ssc-miR-375), and *Drosophila melanogaster* (dme-miR-375-3p). The conserved seed sequences are depicted in red. **B**, Confocal microphotographs (x200) of representative midgut sections of *esg^F/O^> CantonS* and *esg^F/O^>375OE Drosophila* in baseline, unchallenged (UC) conditions. Nuclei are shown in blue (DAPI). Esg+ progenitor cells (GFP+) and their differentiated progeny (GFP+) are shown in green. **C**, Confocal microphotographs (x200) of representative midgut sections of *esg^TS^>CantonS* and *esg^TS^>375OE Drosophila* in baseline, UC conditions. Nuclei are shown in blue (DAPI). Esg+ progenitor cells (GFP+) are shown in green. **D**, Confocal microphotographs (x200) of representative midgut sections of *esg^TS^>CantonS* and *esg^TS^>375OE Drosophila* infected with *Erwinia carotovora carotovora* 15 (*Ecc*15) for 12 h. Nuclei are shown in blue (DAPI). Esg+ progenitor cells (GFP+) are shown in green. Yellow scalebar indicates 50 μm. **E**, Bar plots of total PH3+ cells in the entire midgut of control *D. melanogaster (esg^TS^>CantonS,* n=22-23) and *D. melanogaster* overexpressing miR-375 only in esg+ progenitor cells *(esg^TS^>375OE,* n=23-26) under UC conditions or orally infected with *Ecc*15 for 12 h. *** P < 0.001 by two-tailed Student’s t-test.

In full, these data demonstrate that both genetic deletion (permanent) and pharmacologic suppression (transient) of miR-375 promotes intestinal epithelial growth phenotypes in murine enteroids *ex vivo* and suggest a function for miR-375 in controlling stress-induced proliferation in the *Drosophila* midgut *in vivo*.

### miR-375 control of enteroid growth is mediated in part by regulation of the Yap1 pathway

To identify candidate mechanisms by which miR-375 exerts its effects on intestinal epithelial proliferation, we mined our previously published small RNA-seq and RNA-seq data in Sox9-Low stem cells under diverse conditions (chow diet, high-fat diet, germ-free, and microbially-conventionalized) that variably influence the proliferative state of intestinal crypts (Beyaz et al., 2016; Peck et al., 2017a). Analysis of these data revealed that the expression of miR-375 and the Hippo signaling pathway factor *Yap1* (Yu et al., 2015), a target of miR-375 (Liu et al., 2010), are very strongly inversely correlated (Pearson’s r = −0.808) (**Fig. 6A**). We then treated enteroids established from 375-KO and WT mice with either 2 μM or 5 μM Yap1 inhibitor (verteporfin) (Liu-Chittenden et al., 2012) and compared to the untreated condition (mock). As expected, verteporfin exerted a dose-dependent suppressive effect (mock vs. 2 μM vs. 3 μM) on the Yap1 target gene *Ctgf* (**Fig. S7A**) (Huntoon et al., 2010). Budding efficiency did not appear to be affected by Yap1 inhibition (**Fig. 6B,C**). However, Yap1 inhibition did reduce enteroid survival and, notably, this effect at 2 μM verteporfin was almost completely rescued by the loss of miR-375, but not at 5 μM verteporfin (**Fig. 6D,E**). In effect, miR-375 deficiency compensates for the anti-survival effect of moderate, but not strong, inhibition of Yap1.

**Figure 6.**
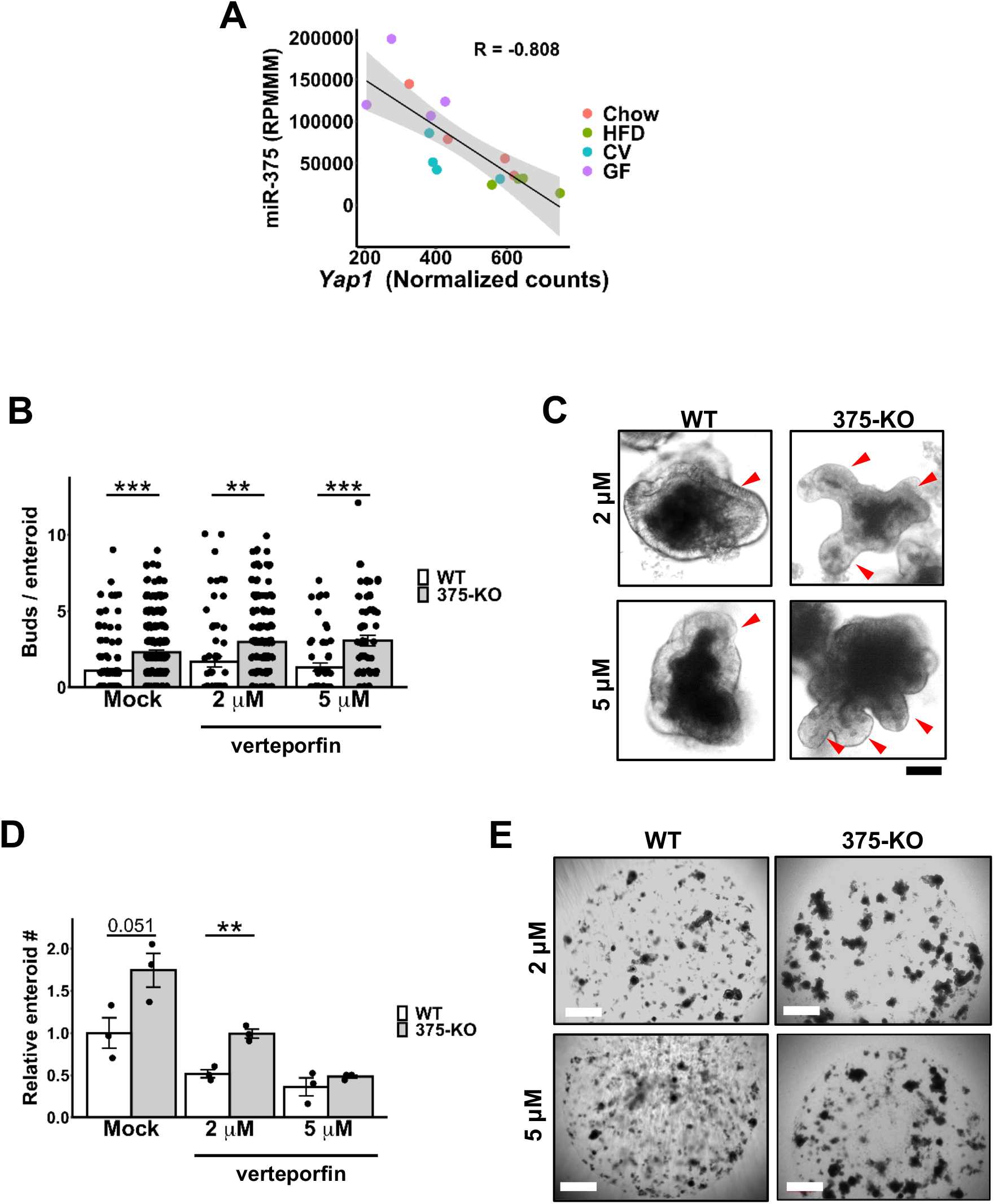
miR-375 influences enteroid growth in part by regulating Yap1 signaling. **A**, Scatter plot of small RNA-seq-determined expression of miR-375 and RNA-seq-determined expression of *Yap1* in jejunal Sox9-Low cells derived from matched mice that were in a diet study in which they were fed either chow (Chow, red) (n=4) or a high fat diet (HFD, green) (n=4) for a prolonged period (16-20 weeks), or were from a study in which the mice were either conventionalized (CV, blue) (n=4) or germfree (GF, purple) (n=4) (Peck, Mah et al.). Pearson correlation analysis revealed a strong inverse correlation (R= −0.808). **B**, Plot of number of buds per enteroid for WT and 375-KO mouse jejunal enteroids that were either mock treated or treated with 2 μM or 5 μM of verteporfin. **C**, Phase-contrast microphotographs (x400) of WT and 375-KO mouse jejunal enteroids treated with either 2 μM or 5 μM of verteporfin. Yellow scalebar indicates 25 μm. **D**, Plot of number of surviving enteroids of WT (n=3) and 375-KO (n=3) mouse jejunal enteroids treated with either 2 μM or 5 μM of verteporfin. **E**, Phase-contrast microphotographs (x50) of WT and 375-KO mouse jejunal enteroids treated with either 2 μM or 5 μM of verteporfin. Yellow scalebar indicates 500 μm. *P < 0.05, **P< 0.01, *** P < 0.001 by two-tailed Student’s t-test. RPMMM, reads per million mapped to microRNAs. RQV, relative quantitative value.

### Intestinal tumors are strongly associated with diminished miR-375 expression

Elevated Wnt (Schneikert and Behrens, 2006) and Yap1 signaling (Zhao et al., 2011) are both associated with the development of intestinal cancers. Therefore, we hypothesized that the development of intestinal tumors may be associated with reduced expression of miR-375. Small intestinal polyps collected from mice with Wnt activating mutations of APC (APCmut), including APCmin and APCq1405x mutations, exhibit decreased levels of miR-375 relative to WT mice (**Fig. S8A**). To determine whether human colon tumors show a similar association, we interrogated the The Cancer Genome Atlas (TCGA) database to compare microRNA levels in primary colon adenocarcinoma tumors to non-tumor colon samples. We found that miR-375 was the most highly expressed miRNA in non-tumor colon tissue that is significantly downregulated (>5-fold) in tumor relative to non-tumor tissue (**Fig. 7A,B**). This pattern of miR-375 expression is preserved when comparing only the matched tumor and non-tumor samples from the same individual (**Fig. 7C**). Notably, integrative analysis with TCGA RNA-seq data revealed that, among miR-375 target genes down-regulated in colon adenocarcinoma samples, *YAP1* is the most inversely correlated gene with miR-375 expression (**Fig. 7D**).

**Figure 7.**
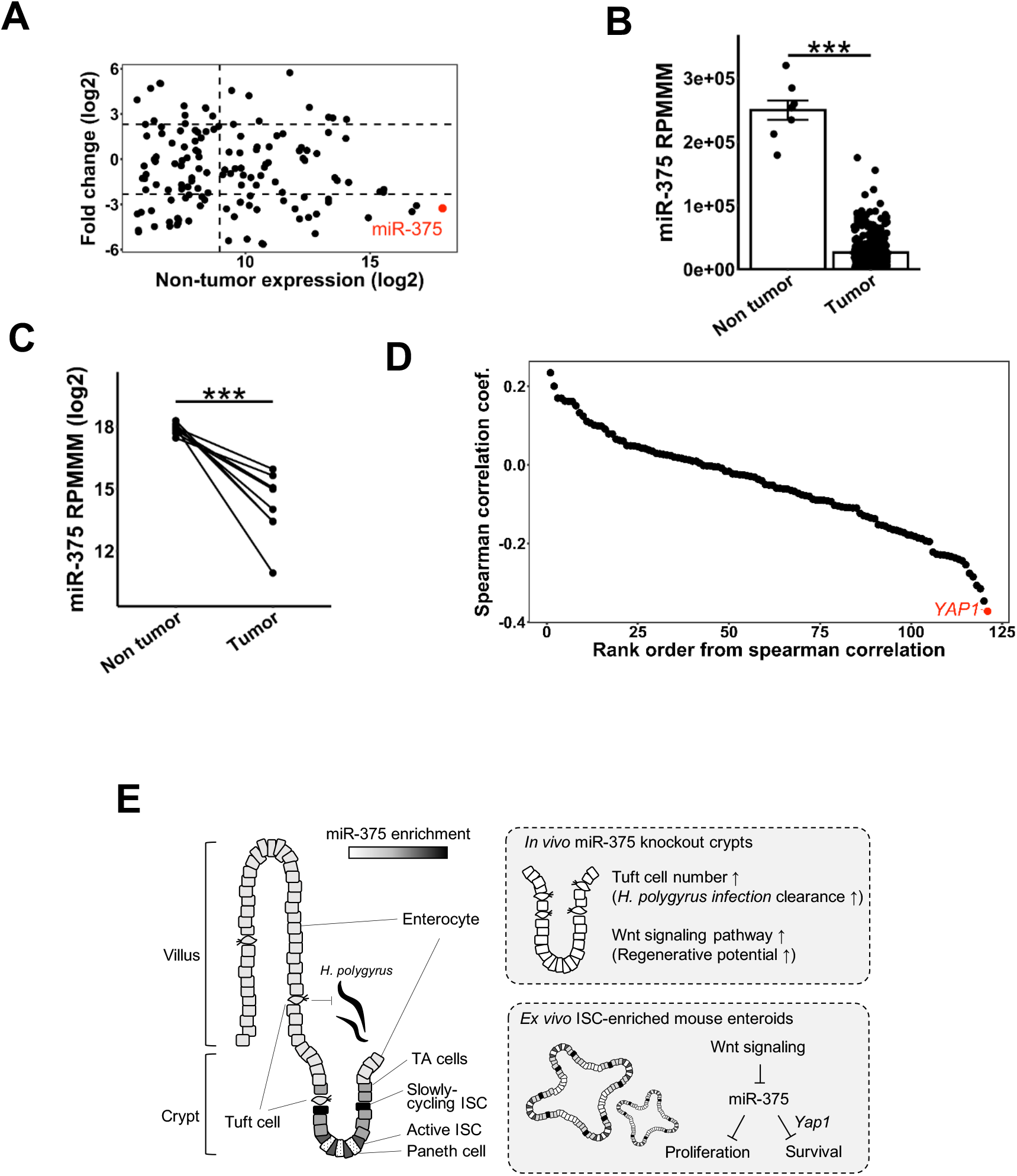
Intestinal cancer is associated with depressed expression of miR-375. **A**, Scatter plot using data from TCGA database to display the fold change of miRNA expression in human primary colon adenocarcinoma (n=371) to human colonic non-tumor tissue (n=8). miR-375 is indicated in red. **B**, Bar plot of miR-375 expression in colon non-tumor tissue (n=8) and primary colon adenocarcinoma (n=371) from TCGA data. **C**, Dot plot of miR-375 expression from TCGA colon non-tumor tissue (n=8) and matched primary, colon adenocarcinoma (n=8) with lines connecting tumors derived from the same patient. **D**, Upregulated genes in TCGA primary colon adenocarcinoma that are predicted miR-375 targets in mouse and human were ranked according to the calculated spearman correlation coefficient to identify inversely correlated genes with miR-375 expression. *YAP1* is indicated in red. **E**, Working model of miR-375 expression and function in the intestinal epithelium. *P < 0.05, *** P < 0.001 by two-tailed Student’s t-test. RQV, relative quantitative value. RPMMM, reads per million mapped to microRNAs.

Taken together, the data presented in this study, which features crypt scRNA-seq, as well as cross-species functional studies, provide a comprehensive view of the role of a miRNA in the intestinal crypt. Important functions of miR-375 revealed in this work include the control of tuft cell abundance and response to helminth *Heligmosomoides polygyrus* infection as well as Wnt-signaling in ISCs and regenerative response to irradiation. A working model of miR-375 expression and function is shown in **Fig. 7E**.

## Discussion

It is recognized that miRNAs affect the structural and functional properties of the small intestinal epithelium (McKenna et al., 2010; Peck et al., 2017a; Singh et al., 2020). Previously, we have shown that an intestinal epithelial cell population that is partially enriched for ISCs is also enriched for several miRNAs that are responsive to microbes (Peck et al., 2017a). In this report, we have: (1) identified 14 miRNAs that are very highly expressed in mouse jejunal ISCs including the most enriched miRNA miR-375; (2) shown that the expression of miR-375 is strongly suppressed by Wnt signaling; (3) found that genetic loss of miR-375 results in an elevation of tuft cell numbers and in the significant reduction of helminth worm burden after *Heligmosomoides polygyrus* infection; (4) determined that miR-375 deficiency promotes the Wnt-signaling pathway in ISCs and enhances the intestinal epithelial regenerative response to irradiation injury; (5) demonstrated that miR-375 regulates intestinal epithelial survival and growth in murine enteroids *ex vivo* and *Drosophila* midgut *in vivo;* (6) shown that the loss of miR-375 can partially compensate for the anti-survival effect of Yap1 inhibition in murine enteroids; and (7) revealed that diminished expression of miR-375 is associated with intestinal tumor development in both mouse and human.

To our knowledge, this is the first report of the profiling of miRNA expression in Lgr5-High ISCs. Additionally, although previous reports from us and others have described the importance of miRNAs in the enteroendocrine cell lineage trajectory (Knudsen et al., 2015; Singh et al., 2020), this study is the first to our knowledge to investigate the function of an ISC-enriched miRNA. Furthermore, this is also the first study to our knowledge that employs single cell RNA-seq to investigate intestinal cell type-specific changes in gene expression in the genetic absence of a miRNA. We found that at baseline the loss of miR-375 promotes tuft cell numbers, and accordingly, enhances eosinophil abundance and reduces *Heligmosomoides polygyrus* burden after infection. The detailed underlying molecular mechanisms merit further investigation. Also, we showed that the loss of miR-375 promotes the intestinal epithelial regenerative response to irradiation. Follow-up functional *ex vivo* studies in enteroids and *in vivo* studies in *Drosophila* midgut revealed that miR-375 contributes to the control of intestinal epithelial cell survival and proliferation, in part by regulation of Yap1. Likely, though, miR-375 has many molecular targets beyond Yap1, and their identification warrant further study in the future.

Luminal changes in microbes and nutrients can also greatly influence the way ISCs effect changes in intestinal epithelial proliferation (Biton et al., 2018; Mah et al., 2014; Peck et al., 2017b). Our earlier report demonstrated that miR-375 is enriched in Sox9-Low cells, which are partially enriched for ISCs, and that its expression is markedly downregulated with exposure to luminal microbes (Peck et al., 2017a). This dependence may mediate the intestinal proliferative changes that are seen during postnatal intestinal development (Al-Nafussi and Wright, 1982) or with exposure to certain antibiotics (Hormann et al., 2014) or certain infectious agents (Santos et al., 2016). In addition to microbial influences, miR-375 has been associated with responding to host metabolic changes. Besides helping to determine the enteroendocrine lineage (Knudsen et al., 2015), it has a well characterized role in regulating secretion of insulin from pancreatic β-cells in response to glucose (Eliasson, 2017). Provision of a chronic high fat diet has been observed to enhance ISC number and the proliferation of the intestinal epithelium (Mah et al., 2014). However, our current examination of the effect of genetic deletion of miR-375 on ISC activity in response to high fat diet has not revealed any significant changes in intestinal epithelial growth properties *in vivo*.

Finally, agents that damage ISCs, such as irradiation and chemotherapies, and that cause acute intestinal injury, will lead to enhanced intestinal proliferation after exposure (Dekaney et al., 2009; Gurley et al., 2017). Stem-like IECs that have lineage determining multipotency, such as those IECs that are enriched in Lower Side Population (LSP) FACS cells, can be induced under these conditions to assume the function of actively cycling ISCs to repopulate and renew the damaged intestinal epithelium (Roche et al., 2015; Yan et al., 2017). It is worth noting that in this report we have found that not only is miR-375 enriched in ISCs relative to enterocytes, it is further enriched in LSP cells relative to Lgr5-High ISCs. It is possible that the elevated expression of miR-375 in these facultative stem cells may help preserve their reserve status by setting a higher threshold for Wnt activation. They may be responsible for the enhancement of intestinal epithelial regeneration that we observe in 375-KO mice in response to radiation injury. Moreover, they may also contribute to the potential Wnt signaling-driven reduction in miR-375 expression that we observe associated with intestinal tumors. Altogether, our data concerning miR-375’s effect on ISC activity does give some indications that miR-375 may be involved in multiple aspects of physiological and pathophysiological regulation of ISCs that merit further detailed investigation in the future.

## Experimental Procedures

### Mouse models

The following mice were utilized: female and male Sox9-EGFP (Formeister et al., 2009), female Lgr5-EGFP (Sato et al., 2009), female and male wild-type C57BL/6, male wild-type and miR-375 null B62J (albino C57BL6/2J), male APCmin, and male APCq1405x. The harvested small intestine was measured and divided into three equal segments. The middle region was considered jejunum. All animal procedures were performed with the approval and authorization of the Institutional Animal Care and Use Committee at each participating institution. Mice were used in these experiments due to their tractability to genetic manipulation, including deletion of the microRNA of interest, as well as the availability of a wide array of appropriate experimental reagents. Mice were housed in well-ventilated cages under 12 hr light/dark cycles with free access to water and standard chow in addition to tubing for environmental enrichment. During experimentation, the mice were monitored at regular intervals to determine their well-being, and at the time of tissue collection the mice were anesthetized by CO2 inhalation and euthanized by means of cervical dislocation.

### Generation of miR-375 knockout mice using CRISPR/Cas9

To delete the miR-375 gene in the mouse, we used the CRISPR/Cas9 system with a guide RNA (gRNA) targeting the 946-965 bp region of the mouse miR-375 gene (ENSMUSG00000065616). 173 FVBxB62J F1 hybrid 1-cell embryos were co-injected with 2.5 μg of gRNA and 7.5 μg of Cas9 mRNA. 107 2-cell embryos were transferred to pseudo-pregnant recipient female mice at ~20 embryos/recipient. 20 founder pups were born and validated for deletion of the miR-375 gene by targeted genomic sequencing. Mice null for miR-375 were backcrossed at least three generations with B62J albino wildtype mice. Synthesis of the gRNA and mouse embryo co-injections and viable embryo emplacement were performed by the Cornell Stem Cell and Transgenic Core Facility at Cornell University (work supported in part by Empire State Stem Cell Fund, contract number C024174).

### Mouse helminth infection

The lifecycle of *Heligmosomoides polygyrus* was maintained in C57BL/6 mice as previously described (Camberis et al., 2003; Johnston et al., 2015). WT and 375-KO B62J mice were infected by oral gavage with 200 *Heligmosomoides polygyrus* L3 larvae. On day 14 post infection, adult worm burdens were assessed by counting the number of worms located in the entire length of the small intestine following exposure of the lumen by dissection.

### Immune cell isolation

For assessment of Type 2 immune cell responses in the intestine, the murine mesenteric lymph nodes (MLNs) were harvested. Single cell suspensions of murine MLNs were prepared by mashing MLNs through a 70 μm cell strainer and counting total number of cells.

### Whole-body mouse irradiation

4-5 months old male WT and 375-KO B62J mice were subjected to 10 Gy of whole-body X-radiation in a cesium-137 irradiator with rotating turntable.

### Mouse high fat diet study

2 months old male WT and 375-KO B62J mice were fed a high fat diet (45% kcal provided by fat) (D12451, Research Diets Inc., New Brunswick, NJ) ad libitum for 14-15 weeks. Body weights were measured weekly.

### Mouse small intestinal villi and polyp isolation

Small intestinal villi were collected from C57BL/6 mice by scraping the mucosal surface of cold PBS-flushed and longitudinally cut whole small intestinal tissue with a glass coverslip. Isolated villi were pelleted by centrifugation (110 x g for 5 minutes at 4°C) and flash frozen. Using a dissection microscope, small intestinal polyps were identified and individually collected from cold PBS-flushed and longitudinally cut small intestine from APCmin and APCq1405x mice. Polyps were subsequently flash frozen.

### Flow cytometry

Four distinct mouse-based cell marker systems were used to sort ISCs (Sox9-EGFP; Lgr5-EGFP; and Cd24) and stem-like IECs (Side Population). Mouse intestinal epithelial cells from the jejunum were dissociated and prepared for fluorescence-activated cell sorting (FACS) as described previously (Mah et al., 2014). For Sox9-EGFP, Lgr5-EGFP, and Side Population cell sorts: CD31-APC (BioLegend, San Diego, CA, cat. 102416), CD45-APC (BioLegend, cat. 1032124), Annexin-V-APC (Life Technologies, Carlsbad, CA, cat. A35110), and Sytox-Blue (Life Technologies, cat. S34857) staining were used to exclude endothelial cells, immune cells, apoptotic cells, and nonviable cells, respectively. The gating parameters of FACS sorting were described previously (Mah et al., 2014). For Cd24-Low cell sorts, UEA-FITC (Vector Laboratories, Burlingame, CA, cat. FL-1061), CD45-FITC (BioLegend, San Diego, CA, cat. 553080), and propidium iodide (BioLegend, cat. 421301) staining was used to exclude goblet/Paneth, endothelial, and nonviable cells, respectively. In addition, for these sorts CD24-Pac Blue (BioLegend, cat. 101819) and EpCAM-PECγ7 (BioLegend, cat. 118215) staining was used to positively select for CD24+ epithelial cells. The Sox9, Lgr5, and Side Population sorts were performed using a Mo-Flo XDP cell sorter (Beckman-Coulter, Fullerton, CA) at the University of North Carolina Flow Cytometry Core Facility. Sorting of Cd24-Low cells was conducted at North Carolina State University, College of Veterinary Medicine using a Mo-Flo XDP cell sorter (Beckman-Coulter, Fullerton, CA). The cells were sorted directly into cold DMEM or lysis buffer.

Side population sorting was used to separate the sub-fraction of slowly cycling from active cycling intestinal stem cells, as described previously (von Furstenberg et al., 2014). Mouse intestinal epithelial cells from the jejunum of female C57BL/6 mice were prepared and sorted into either upper side population (consisting of actively cycling stem cells) or lower side population (consisting of slowly cycling stem cells) by the previously described gating methods (von Furstenberg et al., 2014). The side population sorting was performed using a Mo-Flo XDP cell sorter (Beckman-Coulter, Fullerton, CA) at the University of North Carolina Flow Cytometry Core Facility. Cells were sorted directly into cold lysis buffer (Norgen Biotek, Thorold, ON, Canada).

Immune cell suspensions from helminth-infected mice were incubated with Aqua Live/Dead Fixable Dye (Life Technologies, Grand Island, NY) and fluorochrome-conjugated monoclonal antibodies (mAbs) against mouse CD3 (17A2), CD4 (GK1.5), CD5 (53-7.3), CD11b (M1/70), CD11c (N418), CD19 (eBio1D3), CD25 (PC61.5), CD45 (30-F11), CD45.1 (A20), CD45.2 (104), CD127 (eBioSB/199), CD90.2 (53-2.1), IL-33R (RMST2-2), IL-25R (MUNC33), NK1.1 (PK136), Gata3 (TWAJ) or Siglec-F (E50-2440, BD Biosciences, San Jose, CA). All antibodies from Thermo Fisher unless otherwise noted. Eosinophils, ILC2s and CD4 T helper 2 cells (Th2) were gated as live, CD45^+^SiglecF^+^CD11b^+^; live, CD45^+^lin^-^CD90^+^CD127^+^ST2^+^CD4^-^ and live, CD45^+^lin^+^CD90^+^CD4^+^Gata3^+^, respectively

### Histological analysis

Mouse mid-jejunal tissue was fixed in 4% (v/v) neutral-buffered paraformaldehyde, embedded in paraffin, and cut into 5 μm sections for various staining experiments. Haemotoxylin and eosin (H&E) staining was performed for morphometric analyses (crypt depth). Immunofluorescent staining of PH3 was performed to visualize proliferating cells. Briefly, sections were incubated with primary antibody (rabbit anit-PH3, 1:100 dilution in immunofluorescence buffer,) (Cell Signaling, Danvers, MA, 9701S) overnight at 4° C followed by goat anti-rabbit Alexa fluor 594 secondary antibody (1:400, Invitrogen, Carlsbad, CA, cat. A1102) incubation for 1 hr at room temperature. Hoechst 33342 (1:1000, Invitrogen, cat. C10637) was used to visualize nuclei. Images were captured using a BX53 Olympus scope (Olympus, Center Valley, PA).

### RNA extraction and real-time qPCR

Total RNA was isolated using the Total or Single-cell RNA Purification kit (Norgen Biotek, Thorold, ON, Canada). High Capacity RNA to cDNA kit (Life Technologies, Grand Island, NY) was used for reverse transcription of RNA. TaqMan microRNA Reverse Transcription kit (Life Technologies) was used for reverse transcription of miRNA. Both miRNA and gene expression qPCR were performed using TaqMan assays (Life Technologies) with either TaqMan Universal PCR Master Mix (miRNA qPCR) or TaqMan Gene Expression Master Mix (mRNA qPCR) per the manufacturer’s protocol on a BioRad CFX96 Touch Real Time PCR Detection System (Bio-Rad Laboratories, Richmond, CA). Reactions were performed in triplicate using either U6 (miRNA qPCR) or Rps9 (mouse mRNA qPCR) as the normalizer.

### Small RNA library preparation and sequencing

The small RNA sequencing of cells from the various cell sorts and from enteroids was conducted at Genome Sequencing Facility of Greehey Children’s Cancer Research Institute at University of Texas Health Science Center at San Antonio. Libraries were prepared using the TriLink CleanTag Small RNA Ligation kit (TriLink Biotechnologies, San Diego, CA). Seven to eight libraries were sequenced per lane with single-end 50x on the HiSeq2500 platform. Raw sequencing data is available through GEO accession GSE151088.

### RNA library preparation and sequencing

RNA-sequencing libraries from the Sox9+-EGFP sorts of chow-fed, high fat diet-fed, conventionalized, and germfree C57BL/6J mice were prepared using the Clonetech SMARTer Ultra Low Input library preparation kit combined with Nextera XT DNA sample preparation kit (Illumina) and sequenced with single-end 100 bp on a HiSeq2000 platform at the UNC High Throughput Sequencing Core Facility, as previously described (Peck et al., 2017a). Raw sequencing data is available through GEO accession GSE151088.

### scRNA-seq library preparation and sequencing

Mouse jejunal crypts from 1 year old male WT and 375-KO B62J mice were isolated as previously described (Mah et al., 2014; Peck et al., 2017a). Isolated crypts were resuspended in an ice cold solution of PBS with 0.04% (w/v) bovine serum albumin, and pelleted at 1000 x g at 4°C for 5 minutes. The crypts were subsequently digested with 0.3U/mL dispase I in HBSS at 37°C for 12 minutes with gentle agitation. After stopping dispase I activity with an addition of fetal bovine serum to a final concentration of 10% (v/v), the single cell suspension was filtered, pelleted at 500 x g at 4°C for 5 minutes, washed with cold HBSS, filtered again, and then resuspended in an ice cold solution of PBS with 0.04% (w/v) bovine serum albumin. Prior to submission, single cell suspensions were triturated by pipetting and evaluated for total viable cell number by using trypan blue staining with a TC20 automated cell counter (Bio-Rad Laboratories, Richmond, CA). Single cell RNA sequencing of these samples was performed at the Cornell University Biotechnology Resource Center. Libraries were prepared using the 10X Genomics Chromium preparation kit (10X Genomics, Pleasanton, CA). Raw sequencing data is available through GEO accession GSE151088.

### Bioinformatics analysis

Small RNA-sequencing reads were aligned to the mouse genome (mm9) and quantified using miRquant 2.0 as previously described (Kanke et al., 2016), with the exception that raw miRNA counts were normalized using either RPMMM or DESeq2 (Love et al., 2014) to determine significance. miRNA annotation was performed using miRbase (r18 for mouse). RNA-sequencing reads were mapped to mouse genome release mm10 using STAR (v2.5.3a) (Dobin et al., 2013) and transcript quantification was performed using Salmon (v0.6.0)(Patro et al., 2017). Differential gene expression analysis was accomplished using DESeq2 (Love et al., 2014). Single cell RNA-sequencing was performed using 10x genomics cellranger software (v3.0.1) and aligned to the mouse genome (mm10) to get a cell count matrix. Single cell RNA-seq data was subsequently filtered, clustered, visualized, and analyzed using Seurat (v3.1.5). Clusters were assigned cell types using markers reported by Haber and coworkers (Haber et al., 2017), and unassigned or immune-like clusters were discarded, resulting in 2614 WT and 3080 375-KO B62J mouse cells.

### TCGA analysis

Data <u>Download</u>: RNA-seq data from 382 primary colon tumor samples and 39 solid normal tissue were downloaded from the TCGA database. High Throughput Sequencing (HTSeq) counts data were downloaded using the NIHGDC Data Transfer Tool and normalized using DESeq2. miRNA data from 371 primary colon tumor samples and 8 solid normal tissue were downloaded from the TCGA database. miRNA quantification data using mirbase21 were downloaded using the NIHGDC Data Transfer Tool. <u>Scatterplot graph to narrow down miR-375</u>:miRNAs were filtered for those that had an average expression above 50 RPMMM in the solid normal tissue (n=8). Fold-change was calculated for each miRNA by adding 0.1 RPMMM to all values and calculating the average expression for all tumor samples (n=371) and dividing by the average for all solid normal tissue (n=8). A log2 normalization was then applied to these fold-change values. The average expression for each miRNA and the corresponding log2 foldchange were then graphed as a scatterplot. <u>All tumor vs non-tumor miR-375 Graph</u>: miR-375 expression was extracted for each primary colon tumor (n= 371) and solid normal colon tissue (n=8). Expression of miR-375 was then graphed according to the tissue condition. <u>Matched tumor vs non-tumor miR375 Graph</u>: Datasets representing the patient matched tumor and solid normal tissue (n=8) were identified using the TCGA IDs. Expression for miR-375 was extracted and graphed. Lines connecting data points link patient matched samples. <u>Correlation analysis graph</u>: Significantly upregulated genes in primary colon tumor samples were identified as those genes with an average expression above 1000 normalized counts in tumor (n=39) or non-tumor (n=382), a Benjamini Hochberg adjusted P-value below 0.05, and a log2 fold-change above 0. Spearman correlation coefficients were calculated for each upregulated gene with miR-375 for those samples with both RNA-seq and miRNA data (n=368). Each of the upregulated genes was ranked based on its Spearman correlation coefficient with miR-375. The determined rank was then graphed against the calculated correlation value.

### Mouse Enteroid culture

Jejunal crypts were isolated from 3-5 month old male WT and 375-KO B62J mice as previously described (Peck et al., 2017a). The isolated crypts (Day 0) were grown into Reduced Growth Factor Matrigel (Corning, Corning, NY, cat. 356231). Advanced DMEM/F12 (Gibco, Gaithersburg, MD, cat. 12634-028) supplemented with GlutaMAX (Gibco, cat. 35050-061), Pen/Strep (Gibco, cat. 15140), HEPES (Gibco, cat. 15630-080), N2 supplement (Gibco, cat. 17502-048), 50 ng/mL EGF (R&D Systems, Minneapolis, MN, cat.2028-EG), 100 ug/mL Noggin (PeproTech, Rocky Hill, NJ, cat. 250-38), 250 ng/uL murine R-spondin (R&D Systems, cat. 3474-RS-050), and 10 mM Y27632 (Enzo Life Sciences, Farmingdale, NY, cat. ALX270-333-M025) was added. For miRNA loss-of-function studies, miRCURY LNA Power Inhibitor against mouse miR-375 (mmu-miR-375-3p) (Qiagen, Hilden, Germany, cat.

Y104101397-DFA) or Power Negative Control A (Qiagen, cat. YI00199006-DDA) was added at 500 nM on Day 0 and supplemented at 250 nM on Day 3. Enteroids at Day 5 were harvested for RNA isolation or fixed in 4% (v/v) paraformaldehyde for whole mount staining. For studies inhibiting Wnt and Yap1 function, enteroids were treated with IWP2 (Tocris Bioscience, Bristol, UK, cat. 3533) and verteporfin (Sigma-Aldrich, St. Louis, MO, cat. SML0534-5ML), respectively. Enteroid cultures were exposed to these inhibitors at Day 0 with the initial medium and at Day 3 with the replenishment medium.

### Whole mount enteroids immunostaining and imaging

The fixed mouse enteroids were permeabilized with 0.5% (v/v) Triton X-100/PBS, washed with PBS containing 0.1% (w/v) BSA/0.02% (v/v) TritonX-100/0.05% (v/v) Tween-20 and blocked with 10% (v/v) normal goat serum. Primary antibodies were used to stain PH3 (rabbit anti-Phospho-Histone H3 (Ser10), 1:100, Cell Signaling, Danvers, MA, cat. 9701S). The staining was visualized by fluorescence microscopy with fluorescent-conjugated secondary antibodies (goat anti rabbit Alexa Fluor 594, 1:400, ThermoFisher,Waltham, MA, cat. A-11034). Nuclei were counterstained with Hoechst 33348 dye (1:1000). The immunofluorescent staining was visualized and z-stack bright field images were taken by a ZEISS Axiovert 200M inverted microscope (Zeiss, Jena, Germany).

### CRISPR/Cas9 editing of mouse enteroids

The proximal half of the small intestine was harvested from ~6-week-old C57BL/6 mice. Crypts were isolated and plated in Matrigel. Cells were grown for 3-4 weeks to allow enteroids to form. Enteroids were then transfected with CRISPR base editing tools and guide RNAs to make the APC^Q883^* edit (Han et al., 2017). Cells were then cultured in the absence of Rspo for ~2 weeks to select for Apc mutant cells. Following selection, cells were frozen down.

### Generation and infection of genetically modified lines of Drosophila

Esg-Gal4; UAS-GFP, tub-Gal80^TS^ (Esg^TS^, progenitor specific) (Micchelli and Perrimon, 2006) or Esg^F/O^ (Esg-Gal4, UAS-GFP, tub-Gal80TS; UAS-FLP, actfrtSTOPfrt-Gal4) (Jiang and Edgar, 2009) fruit flies were crossed to UAS lines (BDSC 59916) for creating flies with miR-375 overexpression in Esg stem/progenitor cells. Parental flies were crossed using ~15 female flies and 5 males. They were then transferred during development in a 12:12 hour light/dark 18°C incubator. The parental generation was removed after 5 days in the 18°C incubator to control for fly density of the F1 progeny. Esg^TS^ or Esg^F/O^ flies were crossed to wild type line CantonS (BDSC: 64349) to generate control flies. Parental lines were maintained at room temperature (~23°C) on standard fly medium (50 g baker yeast, 30 g cornmeal, 20 g sucrose, 15 g agar, 5 mL 99% (v/v) propionic acid mix, 0.5 mL 85% (v/v) phosphoric acid, 26.5 mL methyl paraben in 1L ethanol) in a 12:12 hours light/dark cycle. Oral infection of pathogen *Erwinia carotovora ssp. carotovora* 15 (*Ecc*15) was performed as previously described (Buchon et al., 2009). Orally treated flies were incubated at 29°C until dissection for analyses.

### Immunostaining of Drosophila midgut

The excised *Drosophila* midguts were fixed in 4% (v/v) paraformaldehyde and washed with 0.1% (v/v) Triton X-100 in PBS. The samples were then incubated for 1 hr in blocking solution (1% (w/v) bovine serum albumin, 1% (v/v) normal donkey serum, and 0.1% (v/v) Triton X-100 in PBS) followed by overnight primary antibody incubation and 2 hr secondary antibody staining. The primary antibody used in this study was rabbit anti-PH3 (1:000, EMD Millipore, Burlington, MA, cat. 06-570). The secondary antibody used in this study was donkey anti-rabbit-555 (1:2000, Thermo Fisher, Waltham, MA, cat. A-31572). DAPI (1:50000) was used to visualize nuclei. Imaging was performed on a Zeiss LSM 700 fluorescent/ confocal inverted microscope (Zeiss, Jena, Germany).

### Statistics

In most figure panels, quantitative data are reported as an average of biological replicates ± standard error of the mean. In figure panels pertaining to whole mount immunofluorescent staining in enteroids, quantitative data are reported as an average per enteroid from an experiment ± standard error of the mean (enteroids from n=2-5 wells per condition). In all analyses, statistical differences were assessed by two-tailed Student’s t-test with threshold P-value < 0.05, unless otherwise specifically noted.

## Acknowledgments

We gratefully acknowledge the following grants that funded the research work described in this study: American Diabetes Association 1-16-ACE-47 (awarded to P.S.), a Cornell Intercampus Collaborative Seed Grant (awarded to P.S. and L.E.D.), NIH/NIAID R01AI130379 (awarded to E.D.T), NIH/NIAID R21AI153934 (awarded to N.B.), American Cancer Society 131461-RSG-17-202-01-TBG (awarded to L.E.D.), NIH/NCI R01CA222517-01A1 (awarded to L.E.D.), NIH/NIDDK R01DK100508 (awarded to C.D.), NIH P50HD076210 (awarded to J.C.S), a Comparative Medicine and Translational Research Training Program Fellowship T32OD011130 (awarded to B.S.), a SUNY Diversity Fellowship (awarded to J.W.V), and a NYSTEM fellowship (awarded to Y-H.H).

## Author Contributions

Conceptualization, M.T.S., M.K., A.P.S., and P.S.; Methodology, M.T.S., M.K., A.P.S., and P.S.; Formal Analysis, M.K., J.W.V., and W.A.P.; Investigation, M.T.S., M.K., A.P.S., J.W.V., A.J.M., O.O.O., A.B., Y-H.H., B.S., J.C.B., R.L.C., E.G.C, V.D.R., and B.C.E.P.; Resources, C.M.D., S.D., J.C.S., L.E.D., N.B., E.D.T., and P.S.; Writing-Original Draft, M.T.S., M.K., and P.S.; Writing-Review & Editing, M.T.S., M.K., A.P.S., J.W.V., O.O.O., A.B., Y-H.H., C.M.D., L.E.D., N.B., E.D.T., and P.S.; Visualization, M.T.S., M.K., A.P.S., and Y-H.H.; Supervision, P.S.; Funding acquisition, J.W.V., Y-H.H., B.S., C.D., J.C.S., L.E.D., N.B., E.D.T., and P.S..

## References

Al-Nafussi, A.I., and Wright, N.A. (1982). Cell kinetics in the mouse small intestine during immediate postnatal life. Virchows Arch B Cell Pathol Incl Mol Pathol 40, 51–62.

Ayyaz, A., Kumar, S., Sangiorgi, B., Ghoshal, B., Gosio, J., Ouladan, S., Fink, M., Barutcu, S., Trcka, D., Shen, J., et al. (2019). Single-cell transcriptomes of the regenerating intestine reveal a revival stem cell. Nature 569, 121–125.

Barker, N. (2014). Adult intestinal stem cells: critical drivers of epithelial homeostasis and regeneration. Nat Rev Mol Cell Biol 15, 19–33.

Barker, N., van Es, J.H., Kuipers, J., Kujala, P., van den Born, M., Cozijnsen, M., Haegebarth, A., Korving, J., Begthel, H., Peters, P.J., et al. (2007). Identification of stem cells in small intestine and colon by marker gene Lgr5. Nature 449, 1003–1007.

Bartel, D.P. (2018). Metazoan MicroRNAs. Cell 173, 20–51.

Beyaz, S., Mana, M.D., Roper, J., Kedrin, D., Saadatpour, A., Hong, S.J., Bauer-Rowe, K.E., Xifaras, M.E., Akkad, A., Arias, E., et al. (2016). High-fat diet enhances stemness and tumorigenicity of intestinal progenitors. Nature 531, 53–58.

Biton, M., Haber, A.L., Rogel, N., Burgin, G., Beyaz, S., Schnell, A., Ashenberg, O., Su, C.W., Smillie, C., Shekhar, K., et al. (2018). T Helper Cell Cytokines Modulate Intestinal Stem Cell Renewal and Differentiation. Cell 175, 1307–1320 e1322.

Bu, P., Wang, L., Chen, K.Y., Srinivasan, T., Murthy, P.K., Tung, K.L., Varanko, A.K., Chen, H.J., Ai, Y., King, S., et al. (2016). A miR-34a-Numb Feedforward Loop Triggered by Inflammation Regulates Asymmetric Stem Cell Division in Intestine and Colon Cancer. Cell Stem Cell 18, 189–202.

Buchon, N., Broderick, N.A., Kuraishi, T., and Lemaitre, B. (2010). Drosophila EGFR pathway coordinates stem cell proliferation and gut remodeling following infection. BMC Biol 8, 152.

Buchon, N., Broderick, N.A., Poidevin, M., Pradervand, S., and Lemaitre, B. (2009). Drosophila intestinal response to bacterial infection: activation of host defense and stem cell proliferation. Cell Host Microbe 5, 200–211.

Camberis, M., Le Gros, G., and Urban, J., Jr. (2003). Animal model of Nippostrongylus brasiliensis and Heligmosomoides polygyrus. Curr Protoc Immunol Chapter 19, Unit 19 12.

Chen, C.H., Luhur, A., and Sokol, N. (2015). Lin-28 promotes symmetric stem cell division and drives adaptive growth in the adult Drosophila intestine. Development 142, 3478–3487.

Dekaney, C.M., Gulati, A.S., Garrison, A.P., Helmrath, M.A., and Henning, S.J. (2009). Regeneration of intestinal stem/progenitor cells following doxorubicin treatment of mice. Am J Physiol Gastrointest Liver Physiol 297, G461–470.

Dekaney, C.M., Rodriguez, J.M., Graul, M.C., and Henning, S.J. (2005). Isolation and characterization of a putative intestinal stem cell fraction from mouse jejunum. Gastroenterology 129, 1567–1580.

Dobin, A., Davis, C.A., Schlesinger, F., Drenkow, J., Zaleski, C., Jha, S., Batut, P., Chaisson, M., and Gingeras, T.R. (2013). STAR: ultrafast universal RNA-seq aligner. Bioinformatics 29, 15–21.

Dow, L.E., O’Rourke, K.P., Simon, J., Tschaharganeh, D.F., van Es, J.H., Clevers, H., and Lowe, S.W. (2015). Apc Restoration Promotes Cellular Differentiation and Reestablishes Crypt Homeostasis in Colorectal Cancer. Cell 161, 1539–1552.

Eliasson, L. (2017). The small RNA miR-375 - a pancreatic islet abundant miRNA with multiple roles in endocrine beta cell function. Mol Cell Endocrinol 456, 95–101.

Flanagan, D.J., Austin, C.R., Vincan, E., and Phesse, T.J. (2018). Wnt Signalling in Gastrointestinal Epithelial Stem Cells. Genes (Basel) 9.

Formeister, E.J., Sionas, A.L., Lorance, D.K., Barkley, C.L., Lee, G.H., and Magness, S.T. (2009). Distinct SOX9 levels differentially mark stem/progenitor populations and enteroendocrine cells of the small intestine epithelium. Am J Physiol Gastrointest Liver Physiol 296, G1108–1118.

Foronda, D., Weng, R., Verma, P., Chen, Y.W., and Cohen, S.M. (2014). Coordination of insulin and Notch pathway activities by microRNA miR-305 mediates adaptive homeostasis in the intestinal stem cells of the Drosophila gut. Genes Dev 28, 2421–2431.

Gebert, L.F.R., and MacRae, I.J. (2019). Regulation of microRNA function in animals. Nat Rev Mol Cell Biol 20, 21–37.

Gerbe, F., Sidot, E., Smyth, D.J., Ohmoto, M., Matsumoto, I., Dardalhon, V., Cesses, P., Garnier, L., Pouzolles, M., Brulin, B., et al. (2016). Intestinal epithelial tuft cells initiate type 2 mucosal immunity to helminth parasites. Nature 529, 226–230.

Gurley, K.E., Ashley, A.K., Moser, R.D., and Kemp, C.J. (2017). Synergy between Prkdc and Trp53 regulates stem cell proliferation and GI-ARS after irradiation. Cell Death Differ 24, 1853–1860.

Haber, A.L., Biton, M., Rogel, N., Herbst, R.H., Shekhar, K., Smillie, C., Burgin, G., Delorey, T.M., Howitt, M.R., Katz, Y., et al. (2017). A single-cell survey of the small intestinal epithelium. Nature 551, 333–339.

Han, T., Schatoff, E.M., Murphy, C., Zafra, M.P., Wilkinson, J.E., Elemento, O., and Dow, L.E. (2017). R-Spondin chromosome rearrangements drive Wnt-dependent tumour initiation and maintenance in the intestine. Nat Commun 8, 15945.

He, X.C., Zhang, J., Tong, W.G., Tawfik, O., Ross, J., Scoville, D.H., Tian, Q., Zeng, X., He, X., Wiedemann, L.M., et al. (2004). BMP signaling inhibits intestinal stem cell self-renewal through suppression of Wnt-beta-catenin signaling. Nat Genet 36, 1117–1121.

Henning, S.J., and von Furstenberg, R.J. (2016). GI stem cells - new insights into roles in physiology and pathophysiology. J Physiol 594, 4769–4779.

Hormann, N., Brandao, I., Jackel, S., Ens, N., Lillich, M., Walter, U., and Reinhardt, C. (2014). Gut microbial colonization orchestrates TLR2 expression, signaling and epithelial proliferation in the small intestinal mucosa. PLoS One 9, e113080.

Huntoon, C.J., Nye, M.D., Geng, L., Peterson, K.L., Flatten, K.S., Haluska, P., Kaufmann, S.H., and Karnitz, L.M. (2010). Heat shock protein 90 inhibition depletes LATS1 and LATS2, two regulators of the mammalian hippo tumor suppressor pathway. Cancer Res 70, 8642–8650.

Ivey, K.N., and Srivastava, D. (2010). MicroRNAs as regulators of differentiation and cell fate decisions. Cell Stem Cell 7, 36–41.

Jiang, H., and Edgar, B.A. (2009). EGFR signaling regulates the proliferation of Drosophila adult midgut progenitors. Development 136, 483–493.

Jiang, L., and Hermeking, H. (2017). miR-34a and miR-34b/c Suppress Intestinal Tumorigenesis. Cancer Res 77, 2746–2758.

Johnston, C.J., Robertson, E., Harcus, Y., Grainger, J.R., Coakley, G., Smyth, D.J., McSorley, H.J., and Maizels, R. (2015). Cultivation of Heligmosomoides polygyrus: an immunomodulatory nematode parasite and its secreted products. J Vis Exp, e52412.

Kanke, M., Baran-Gale, J., Villanueva, J., and Sethupathy, P. (2016). miRquant 2.0: an Expanded Tool for Accurate Annotation and Quantification of MicroRNAs and their isomiRs from Small RNA-Sequencing Data. J Integr Bioinform 13, 307.

Kim, K., Hung, R.J., and Perrimon, N. (2017). miR-263a Regulates ENaC to Maintain Osmotic and Intestinal Stem Cell Homeostasis in Drosophila. Dev Cell 40, 23–36.

Knudsen, L.A., Petersen, N., Schwartz, T.W., and Egerod, K.L. (2015). The MicroRNA Repertoire in Enteroendocrine Cells: Identification of miR-375 as a Potential Regulator of the Enteroendocrine Lineage. Endocrinology 156, 3971–3983.

Liu-Chittenden, Y., Huang, B., Shim, J.S., Chen, Q., Lee, S.J., Anders, R.A., Liu, J.O., and Pan, D. (2012). Genetic and pharmacological disruption of the TEAD-YAP complex suppresses the oncogenic activity of YAP. Genes Dev 26, 1300–1305.

Liu, A.M., Poon, R.T., and Luk, J.M. (2010). MicroRNA-375 targets Hippo-signaling effector YAP in liver cancer and inhibits tumor properties. Biochem Biophys Res Commun 394, 623–627.

Love, M.I., Huber, W., and Anders, S. (2014). Moderated estimation of fold change and dispersion for RNA-seq data with DESeq2. Genome Biol 15, 550.

Mah, A.T., Van Landeghem, L., Gavin, H.E., Magness, S.T., and Lund, P.K. (2014). Impact of diet-induced obesity on intestinal stem cells: hyperproliferation but impaired intrinsic function that requires insulin/IGF1. Endocrinology 155, 3302–3314.

McKenna, L.B., Schug, J., Vourekas, A., McKenna, J.B., Bramswig, N.C., Friedman, J.R., and Kaestner, K.H. (2010). MicroRNAs control intestinal epithelial differentiation, architecture, and barrier function. Gastroenterology 139, 1654–1664, 1664 e1651.

Micchelli, C.A., and Perrimon, N. (2006). Evidence that stem cells reside in the adult Drosophila midgut epithelium. Nature 439, 475–479.

Mo, M.L., Li, M.R., Chen, Z., Liu, X.W., Sheng, Q., and Zhou, H.M. (2013). Inhibition of the Wnt palmitoyltransferase porcupine suppresses cell growth and downregulates the Wnt/beta-catenin pathway in gastric cancer. Oncol Lett 5, 1719–1723.

Nalapareddy, K., Nattamai, K.J., Kumar, R.S., Karns, R., Wikenheiser-Brokamp, K.A., Sampson, L.L., Mahe, M.M., Sundaram, N., Yacyshyn, M.B., Yacyshyn, B. et al. (2017). Canonical Wnt Signaling Ameliorates Aging of Intestinal Stem Cells. Cell Rep 18, 2608–2621.

Patro, R., Duggal, G., Love, M.I., Irizarry, R.A., and Kingsford, C. (2017). Salmon provides fast and bias-aware quantification of transcript expression. Nat Methods 14, 417–419.

Peck, B.C., Mah, A.T., Pitman, W.A., Ding, S., Lund, P.K., and Sethupathy, P. (2017a). Functional Transcriptomics in Diverse Intestinal Epithelial Cell Types Reveals Robust MicroRNA Sensitivity in Intestinal Stem Cells to Microbial Status. J Biol Chem 292, 2586–2600.

Peck, B.C.E., Shanahan, M.T., Singh, A.P., and Sethupathy, P. (2017b). Gut Microbial Influences on the Mammalian Intestinal Stem Cell Niche. Stem Cells Int 2017, 5604727.

Peng, S., Gao, D., Gao, C., Wei, P., Niu, M., and Shuai, C. (2016). MicroRNAs regulate signaling pathways in osteogenic differentiation of mesenchymal stem cells (Review). Mol Med Rep 14, 623–629.

Richmond, C.A., Shah, M.S., Carlone, D.L., and Breault, D.T. (2016). An enduring role for quiescent stem cells. Dev Dyn 245, 718–726.

Roche, K.C., Gracz, A.D., Liu, X.F., Newton, V., Akiyama, H., and Magness, S.T. (2015). SOX9 maintains reserve stem cells and preserves radioresistance in mouse small intestine. Gastroenterology 149, 1553–1563 e1510.

Santos, A.J.M., Durkin, C.H., Helaine, S., Boucrot, E., and Holden, D.W. (2016). Clustered Intracellular Salmonella enterica Serovar Typhimurium Blocks Host Cell Cytokinesis. Infect Immun 84, 2149–2158.

Sato, T., Vries, R.G., Snippert, H.J., van de Wetering, M., Barker, N., Stange, D.E., van Es, J.H., Abo, A., Kujala, P., Peters, P.J., et al. (2009). Single Lgr5 stem cells build crypt-villus structures in vitro without a mesenchymal niche. Nature 459, 262–265.

Schneikert, J., and Behrens, J. (2006). Truncated APC is required for cell proliferation and DNA replication. Int J Cancer 119, 74–79.

Singh, A.P., Hung, Y.H., Shanahan, M.T., Kanke, M., Bonfini, A., Dame, M.K., Biraud, M., Peck, B.C.E., Oyesola, O.O., Freund, J.M., et al. (2020). Enteroendocrine Progenitor Cell-Enriched miR-7 Regulates Intestinal Epithelial Proliferation in an Xiap-Dependent Manner. Cell Mol Gastroenterol Hepatol 9, 447–464.

Tao, S., Tang, D., Morita, Y., Sperka, T., Omrani, O., Lechel, A., Sakk, V., Kraus, J., Kestler, H.A., Kuhl, M., et al. (2015). Wnt activity and basal niche position sensitize intestinal stem and progenitor cells to DNA damage. EMBO J 34, 624–640.

Tian, Y., Ma, X., Lv, C., Sheng, X., Li, X., Zhao, R., Song, Y., Andl, T., Plikus, M.V., Sun, J., et al. (2017). Stress responsive miR-31 is a major modulator of mouse intestinal stem cells during regeneration and tumorigenesis. Elife 6.

von Furstenberg, R.J., Buczacki, S.J., Smith, B.J., Seiler, K.M., Winton, D.J., and Henning, S.J. (2014). Side population sorting separates subfractions of cycling and non-cycling intestinal stem cells. Stem Cell Res 12, 364–375.

von Furstenberg, R.J., Gulati, A.S., Baxi, A., Doherty, J.M., Stappenbeck, T.S., Gracz, A.D., Magness, S.T., and Henning, S.J. (2011). Sorting mouse jejunal epithelial cells with CD24 yields a population with characteristics of intestinal stem cells. Am J Physiol Gastrointest Liver Physiol 300, G409–417.

Yan, K.S., Gevaert, O., Zheng, G.X.Y., Anchang, B., Probert, C.S., Larkin, K.A., Davies, P.S., Cheng, Z.F., Kaddis, J.S., Han, A., et al. (2017). Intestinal Enteroendocrine Lineage Cells Possess Homeostatic and Injury-Inducible Stem Cell Activity. Cell Stem Cell 21, 78–90 e76.

Yu, F.X., Meng, Z., Plouffe, S.W., and Guan, K.L. (2015). Hippo pathway regulation of gastrointestinal tissues. Annu Rev Physiol 77, 201–227.

Zhang, H., Wang, Y., Yang, G., Yu, H., Zhou, Z., and Tang, M. (2019). MicroRNA-30a regulates chondrogenic differentiation of human bone marrow-derived mesenchymal stem cells through targeting Sox9. Exp Ther Med 18, 4689–4697.

Zhao, B., Tumaneng, K., and Guan, K.L. (2011). The Hippo pathway in organ size control, tissue regeneration and stem cell self-renewal. Nat Cell Biol 13, 877–883.

