## Supplement Figures for "Single Cell Analysis Reveals Multi-faceted miR-375 Regulation of the Intestinal Crypt"

#### Supplemental Figure Legends

**Figure Supplement 1. Validation of isolated intestinal crypts, villi, and FACS sorted jejunal IECs. A**, RT-qPCR data of *Lgr5* expression in jejunal villi relative to crypts in WT mice (n=4). **B**, RT-qPCR data of *Sis* expression in jejunal villi relative to crypts in WT mice (n=4). **C**, RT-qPCR data of *Lgr5*, *Muc2* and *ChgA* expression in jejunal Lgr5-High cells (n=2) relative to jejunal Lgr5-Low+Neg cells (n=2). **D**, RT-qPCR data of *Lgr5*, *Sis*, *Muc2*, *ChgA*, *Lyz1*, *Dclk1*, and *Bmi1* expression in jejunal Sox9-Neg cells (n=3) and jejunal Sox9-Low cells (n=3) relative to jejunal unsorted cells (n=3). **E**, RT-qPCR data for *Lgr5*, *Muc2* and *ChgA* expression in jejunal Upper Side Population (USP) (n=3) and jejunal Lower Side Population (LSP) cells (n=3) relative to jejunal non-Side Population sorted cells (n=3). *P < 0.05, **P< 0.01 by two-tailed Student's t-test. RQV, relative quantitative value.

**Figure Supplement 2. Validation of the genotype of mice. A**, DNA agarose gel electrophoretogram of mouse tail DNA of 5 months old and 1year old WT and 375-KO B62J mice PCR amplified for the miR-375 gene. A 100bp ladder was used for amplicon size determination. **B**, RT-qPCR data of miR-375 expression of mouse jejunal enteroids of 375-KO (n=3) mice relative to WT (n=3). *** P < 0.001 by two-tailed Student's t-test. RQV, relative quantitative value.

**Figure Supplement 3. Helminth *Heligmosomoides polygyrus* infection of miR-375 null mice does not alter the number of total immune cells. A**, Bar plot of total small intestinal immune cells from FACS analyzed small intestinal tissue from helminth *Heligmosomoides polygyrus* infected WT (n=10) and 375-KO (n=10) B62J mice after 14 days post-inoculation.

**Figure Supplement 4. miR-375 does not affect intestinal architecture at baseline or under chronic high fat diet.** **A**, Plot of crypt depth of the mid-jejunum of 5 months old WT (n=4) and 375-KO (n=8) B62J mice. **B**, Representative H&E stained brightfield microphotographs (x200) of the mid-jejunum of 5 months old WT and 375-KO mice. Yellow scalebar indicates 40 µm. **C**, Plot of number of PH3+ cells per mid-jejunal crypt of 5 months old 375-KO mice (n=4) relative to WT mice (n=4). **D**, Fluorescent photomicrographs (x200) of the mid-jejunum of 5 months old WT and 375-KO mice. Hoechst-stained nuclei are shown in blue and PH3+ cells are shown in red. Yellow scalebar indicates 40 µm. **E**, Percentage weight gain of WT (n=4) and 375-KO (n=4) mice at weekly intervals over 16 weeks of a high fat diet. **F**, Plot of crypt depth of the mid-jejunum of 5 months old WT (n=4) and 375-KO (n=4) mice that were subject to 15 weeks of a high fat diet. **G**, Representative H&E stained brightfield microphotographs (x200) of the mid-jejunum of high fat diet fed 5 months old WT and 375-KO mice. Yellow scalebar indicates 40 µm.

**Figure Supplement 5. Whole-body irradiation of WT albino mice.** **A**, Representative H&E stained brightfield microphotographs (x100) of the mid-jejunum of 5 months old WT B62J mice that were non-irradiated or irradiated for 1 or 2.5 days. Yellow scalebar indicates 50 µm. A, RT-qPCR data of expression of the Yap1 target gene *Ctgf* in wildtype (WT) B62J or 375-KO mouse jejunal enteroids under mock conditions (n=2) or treated with 2 µM (n=2) or 3 µM (n=2) of the Yap1 inhibitor verteporfin.

**Figure Supplement 6. Validation of in vitro knockdown of miR-375 expression in enteroids.** **A**, RT-qPCR data of miR-375 expression from WT B62J jejunal enteroids that were mock treated (n=5) or treated with LNA-scr (n=5) or LNA-375 (n=5). **B**, Plot of enteroid size of mock (n=39) , LNA-scr, (n=32) and LNA-375 (n=56) treated WT B62J mouse jejunal enteroids. **C**, Phase-contrast microphotographs (x400) of WT B62J mouse jejunal enteroids that were either mock, LNA-scr, or LNA-375 treated. White outlines in the images illustrate the measured enteroid areas. Yellow scalebar indicates 50 μm. *** P < 0.001 by two-tailed Student's t-test. RQV, relative quantitative value. B, RT-qPCR data of expression of the Yap1 target gene *Ctgf* in wildtype (WT) B62J or 375-KO mouse jejunal enteroids under mock conditions (n=2) or treated with 2 μM (n=2) or 3 μM (n=2) of the Yap1 inhibitor verteporfin.

**Figure Supplement 7. Validation of verteporfin-mediated inhibition of Yap1 signaling.** **A**, RT-qPCR data of expression of the Yap1 target gene *Ctgf* in wildtype (WT) B62J or 375-KO mouse jejunal enteroids under mock conditions (n=2) or treated with 2 μM (n=2) or 3 μM (n=2) of the Yap1 inhibitor verteporfin.

**Figure Supplement 8. miR-375 expression in APC deficient mouse-derived small intestinal polyps.** **A**, Bar plot of miR-375 expression of C57BL/6 wildtype (WT) villi (n=2) and small intestinal polyps derived from mice harboring inactivating APC mutations (APCmut) (n=6), including APCmin and APCq1405x mutations.

**Table Supplement 1.** Significant differentially expressed microRNAs of jejunal crypts relative to jejunal villi of 3-5 months old WT B62J mice with associated log2 fold-change values and P-values. MicroRNAs are ranked by their fold-change values.

**Table Supplement 2.** Comparative listing of enriched microRNAs in Sox9-Low, Lgr5-High and Cd24-Low cells. MicroRNAs are ranked by expression level. Red labeled microRNAs are enriched in all three cell types.

**Table Supplement 3.** Significant differentially expressed microRNAs of jejunal APC-mut mouse enteroids relative to jejunal wildtype C57BL/6 wildtype (WT) mouse enteroids with associated log2 fold-change values and adjusted P-values. MicroRNAs are ranked by their fold-change values.

### Figure S1

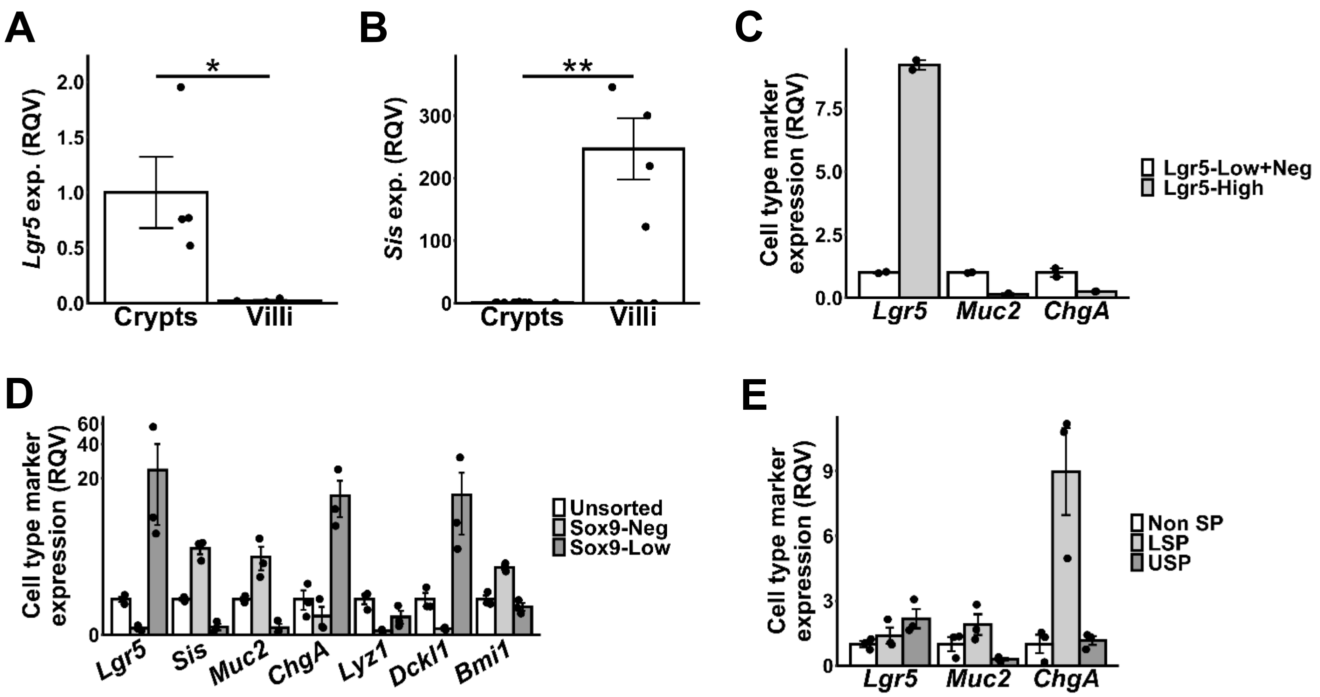
A
Lgr5 exp. (RQV)
2.0
1.5
1.0
0.5
0.0
*
Crypts
Villi
B
Sis exp. (RQV)
300
200
100
0
**
Crypts
Villi
C
Cell type marker expression (RQV)
7.5
5.0
2.5
0.0
Lgr5
Muc2
ChgA
Lgr5-Low+Neg
Lgr5-High
D
Cell type marker expression (RQV)
60
40
20
0
Lgr5
Sis
Muc2
ChgA
Lyz1
Dckl1
Bmi1
Unsorted
Sox9-Neg
Sox9-Low
E
Cell type marker expression (RQV)
9
6
3
0
Lgr5
Muc2
ChgA
Non SP
LSP
USP

Figure S2

A

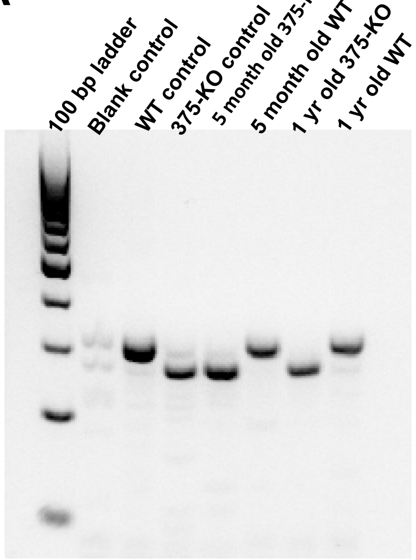
100 bp ladder
Blank control
WT control
375-KO control
5 month old 375-KO
5 month old WT
1 yr old 375-KO
1 yr old WT

B

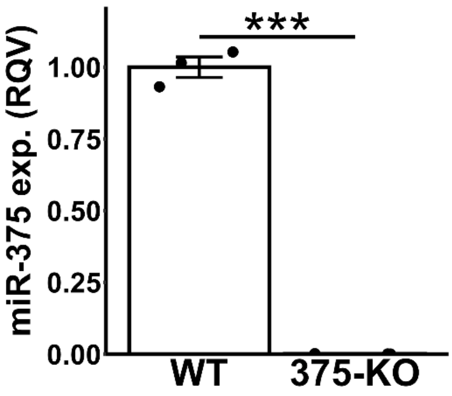
miR-375 exp. (RQV)
1.00
0.75
0.50
0.25
0.00
***
WT
375-KO

### Figure S3

A

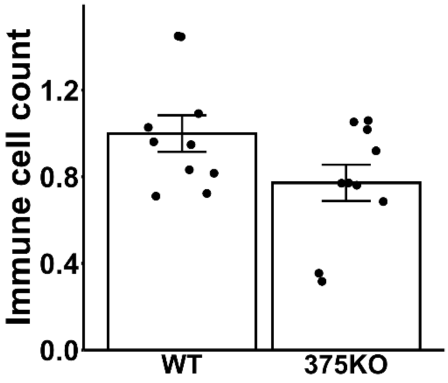

### Figure S4

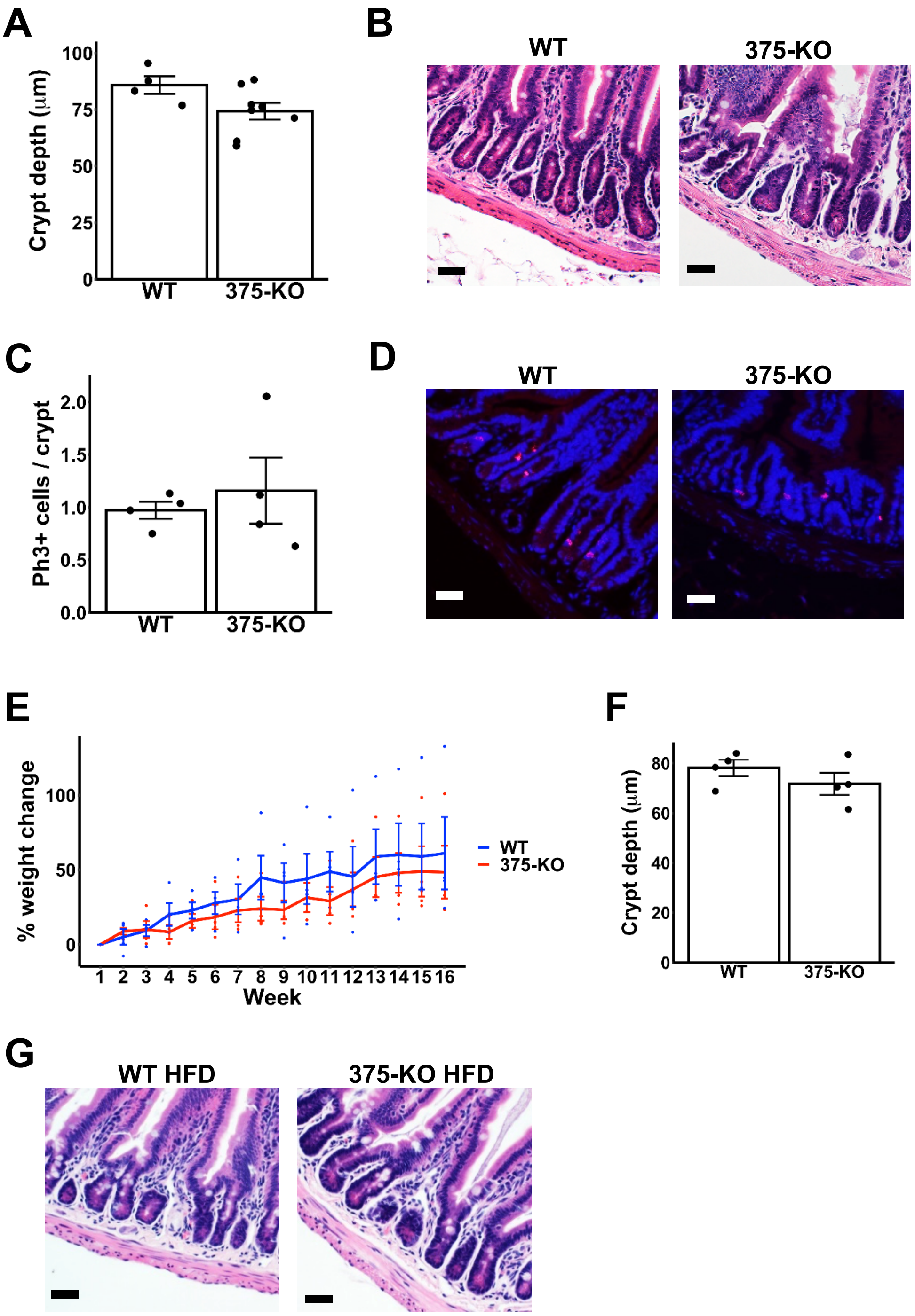

A
Crypt depth (μm)
100
75
50
25
0
WT
375-KO
B
WT
375-KO
C
Ph3+ cells / crypt
2.0
1.5
1.0
0.5
0.0
WT
375-KO
D
WT
375-KO
E
% weight change
100
50
0
1 2 3 4 5 6 7 8 9 10 11 12 13 14 15 16
Week
WT
375-KO
F
Crypt depth (μm)
80
60
40
20
0
WT
375-KO
G
WT HFD
375-KO HFD

### Figure S5

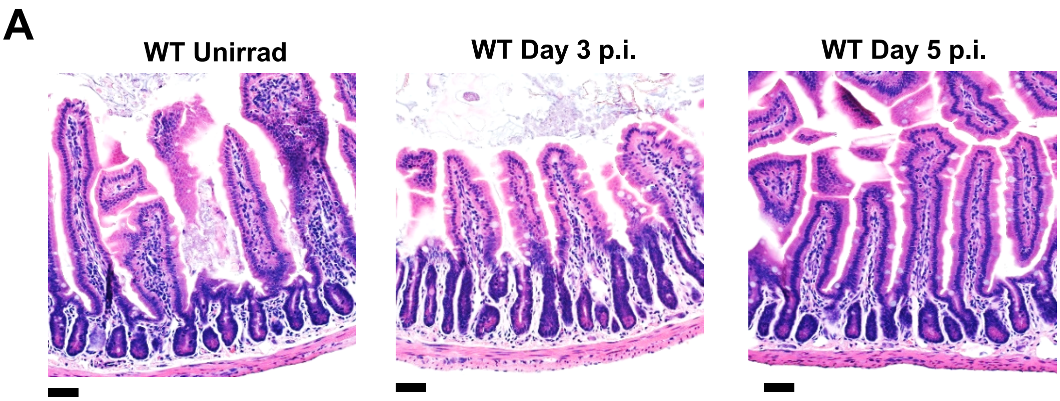

### Figure S6

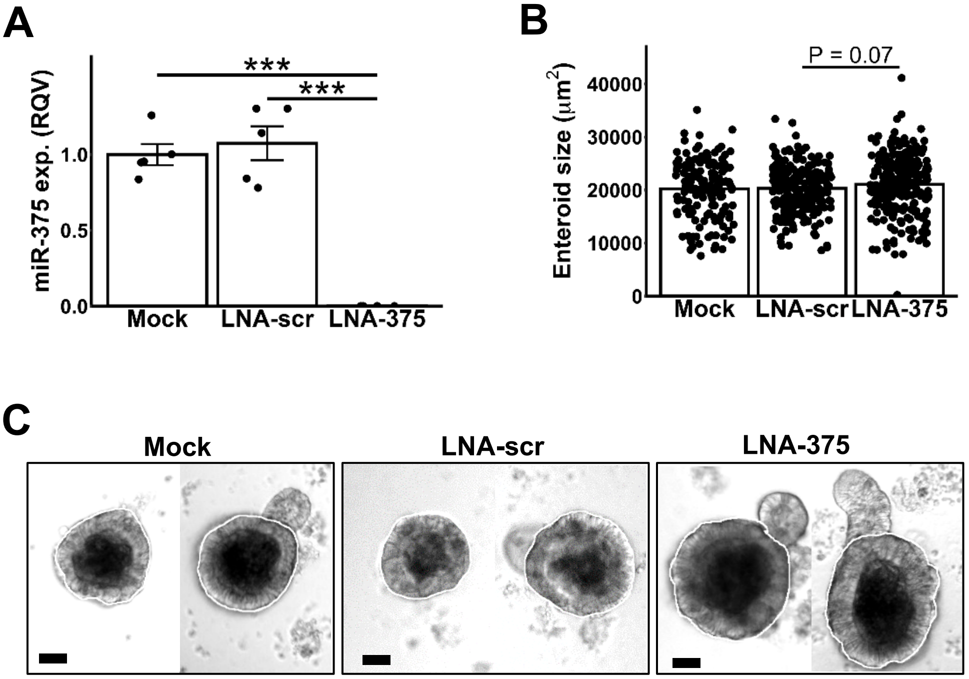
A
miR-375 exp. (RQV)
0.0
0.5
1.0
***
***
Mock
LNA-scr
LNA-375
B
Enteroid size (μm²)
0
10000
20000
30000
40000
P = 0.07
Mock
LNA-scr
LNA-375
C
Mock
LNA-scr
LNA-375

### Figure S7

**A**

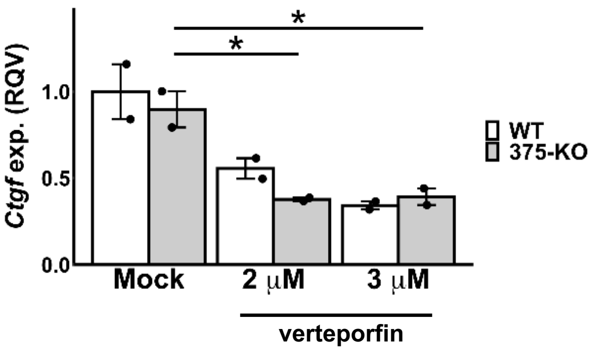

### Figure S8

A

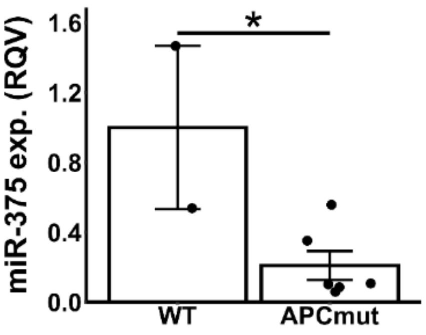

### Table S1

| miR | log2(fold change) | p-value |
|---|---|---|
| miR-155-5p | 2.32 | 0.00571 |
| miR-328-3p | 1.61 | 6.90E-04 |
| miR-92a-1-3p | 1.45 | 6.50E-04 |
| miR-375-3p | 1.43 | 0.00274 |
| let-7e-5p | 1.41 | 0.0091 |
| miR-5099-2 | 1.38 | 8.50E-04 |
| miR-181a-1-5p | 1.33 | 1.70E-04 |
| miR-181a-2-5p | 1.33 | 1.70E-04 |
| miR-194-2-5p | -1.37 | 7.90E-04 |
| miR-194-1-3p | -1.49 | 0.0214 |
| miR-215-3p | -1.57 | 0.00108 |
| miR-194-1-5p | -1.78 | 1.00E-04 |
| miR-215-5p | -2.18 | 8.50E-04 |

### Table S2

| Sox9-Low | Lgr5-High | Cd24-Low |
|---|---|---|
| miR-192-5p | miR-192-5p | miR-215-5p |
| let-7b-5p | let-7b-5p | miR-192-5p |
| miR-375-3p | miR-215-5p | let-7b-5p |
| let-7c-2-5p | let-7c-2-5p | miR-21-5p |
| let-7c-1-5p | let-7c-1-5p | miR-194-1-5p |
| miR-215-5p | miR-192-5p+1 | miR-375-3p |
| miR-92a-1-3p | let-7a-1-5p | miR-192-5p+1 |
| miR-7a-1-5p | let-7a-2-5p | let-7c-2-5p |
| miR-7a-2-5p | miR-200c-3p | let-7c-1-5p |
| miR-191-5p | let-7f-2-5p | miR-26a-2-5p |
| miR-26a-2-5p | miR-375-3p | miR-26a-1-5p |
| miR-26a-1-5p | let-7d-5p | let-7f-2-5p |
| miR-192-5p+1 | let-7f-1-5p | let-7f-1-5p |
| let-7i-5p | miR-191-5p | miR-200c-3p |
| miR-200a-3p | miR-200a-3p | miR-378-3p |
| let-7f-2-5p | miR-26a-2-5p | miR-7a-1-5p |
| miR-21-5p | miR-26a-1-5p | let-7g-5p |
| let-7f-1-5p | miR-21-5p | miR-7a-2-5p |
| let-7a-1-5p | miR-141-3p | miR-200a-3p |
| let-7a-2-5p | miR-182-5p | let-7a-1-5p |

Red denotes miRs present in all three, columns ordered by expression level

### Table S3

| miR | log2(fold change) | adjusted p-value |
|---|---|---|
| miR-146a-5p | 5.99 | 6.04461E-06 |
| miR-10a-5p+1 | 3.45 | 2.40E-03 |
| miR-10a-5p | 3.37 | 3.11E-03 |
| miR-99b-5p | 2.60 | 5.85201E-07 |
| let-7e-5p | 2.49 | 1.48273E-08 |
| miR-125a-5p | 2.25 | 4.07E-04 |
| miR-375-3p | -1.69 | 9.00E-04 |
| miR-141-3p | -1.82 | 5.48E-04 |
| miR-192-5p+1 | -1.83 | 4.11E-04 |
| miR-192-5p | -2.00 | 0.00022 |
| miR-802-5p | -2.15 | 0.0016 |
| miR-22-3p | -2.21 | 2.48E-19 |
| miR-194-5p | -3.59 | 1.68E-10 |
| miR-218-5p | -3.77 | 0.0019 |
